# Modulation of transcription burst amplitude underpins dosage compensation in the *Drosophila* embryo

**DOI:** 10.1101/2023.02.03.526973

**Authors:** Lauren Forbes Beadle, Hongpeng Zhou, Magnus Rattray, Hilary L. Ashe

## Abstract

Dosage compensation, the balancing of X linked gene expression between sexes and to the autosomes, is critical to an organism’s fitness and survival. In *Drosophila*, dosage compensation involves hypertranscription of the male X chromosome. Here we use quantitative live imaging and modelling at single-cell resolution to determine the mechanism underlying X chromosome dosage compensation in *Drosophila*. We show that the four X chromosome genes studied undergo transcriptional bursting in male and female embryos. Mechanistically our data reveal that transcriptional upregulation of male X chromosome genes is primarily mediated by a higher RNA polymerase II initiation rate and burst amplitude across the expression domain. In contrast, burst frequency is spatially modulated in nuclei within the expression domain in response to different transcription factor concentrations to tune the transcriptional response. Together, these data show how the local and global regulation of distinct burst parameters establish the complex transcriptional outputs underpinning developmental patterning.

## Introduction

Dosage compensation was originally discovered in *Drosophila*^1^, where it was found that males increase the transcription of most active X chromosome genes up to 2-fold.^2–4^ In *Drosophila*, the most widely supported models of dosage compensation include a direct role for the male specific lethal (MSL) complex, which targets the male X chromosome. This complex is composed of five proteins - MSL1-3, Maleless and the males absent on the first (MOF) histone acetyltransferase - and two non-coding RNAs transcribed from the X chromosome, *RNA on the X* (*roX*)*1* and *roX2*. Dosage compensation is restricted to males as, based on an X:autosome ratio of 1, the Sex lethal RNA binding protein accumulates in female embryos and represses the translation of *msl2* mRNAs.^5,6^

A favoured model for targeting of the MSL complex is that it binds to high affinity sites (HASs) on the X chromosome, which include *roX1* and r*oX2*^7,8^, then spreads along the X chromosome to the bodies of active genes.^7–11^ Recruitment of the MSL complex to these HASs requires the CLAMP transcription factor.^12^ In the early embryo CLAMP initially binds genome-wide and recruits the MSL complex, before both become enriched at HASs on the male X chromosome.^13^ However, recently an alternative model has been proposed whereby MSL2 and the *roX* RNAs trap the MSL complex on the X chromosome to nucleate a compartment that is necessary for dosage compensation. Evidence for this model includes the finding that the *roX* RNAs and MSL2, via its intrinsically disordered C terminal domain, form stable condensates.^14^

An alternative model for dosage compensation, the inverse dosage model, also exists. This model is based on the idea of genomic balance from studies of aneuploidy and polyploidy, where there is a negative correlation between gene expression and chromosomal dosage.^15^ It posits that the single X chromosome in males results in altered stoichiometry and activity of multi-subunit complexes, such as those involved in gene regulation, which would result in an upregulation of the entire genome.^16,17^ In this model, the MSL complex is not directly required for X chromosome transcriptional upregulation. Instead MSL targeting to the X sequesters MOF and other histone modifiers away from the autosomes to mute their transcriptional upregulation. An additional activity is also suggested to constrain X chromosome transcription that could arise from the high levels of histone acetylation due to MOF.^15^

An RNA polymerase II (Pol II) elongation-based mechanism has been proposed to explain the doubling of transcription on the male X chromosome. This ‘jump start and gain’ model is based on nascent RNA sequencing and Pol II ChIP-chip data from tissue culture cells, which suggested an MSL complex-dependent enhancement of Pol II levels at the 3’ ends of gene bodies. Elevated elongation was postulated to be a result of enhanced release of Pol II from 5’ pausing, the ‘jump’, and improved Pol II processivity, the ‘gain’.^18,19^ As MOF within the MSL complex acetylates H4K16 predominantly on the X chromosome^20–23^, this modification was proposed to reduce the steric hindrance of nucleosomes to Pol II.^18,19^ However, an alternative initiation mechanism due to increased Pol II recruitment has also been proposed, based on a comparison of Pol II ChIP-seq data from male, female and MSL2 knockdown male salivary glands. Higher Pol II was found at the promoters of a subset of genes on the wildtype male X^24^, although the relevance of the small (~1.2 fold) change in promoter Pol II levels has been questioned.^25–27^

Advances in imaging have revealed that many genes are transcribed in discontinuous bursts of transcriptional activity, in organisms ranging from bacteria to mammals.^28^ In this study, we exploit live and quantitative imaging to determine whether hypertranscription of male X chromosome genes involves a higher frequency and/or amplitude of transcriptional bursts. Our data suggest that dosage compensation is mediated by a higher amplitude of transcriptional bursts in male embryos. In contrast, burst frequency is tuned locally in a sex-independent manner to coordinate the transcriptional output to local transcription factor inputs.

## Results

### Live imaging of dosage compensated transcription in the *Drosophila* embryo

To investigate dosage compensation in the early *Drosophila* embryo we chose four X chromosome genes, *short gastrulation (sog*), *hindsight (hnt*), *giant (gt*) and *multiple edematous wings* (*mew*). These genes were chosen based on published time series RNA-seq data from male and female embryos, which found that *sog*, *gt* and *hnt* are compensated in the early embryo, whereas *mew* is not effectively compensated.^29^ To investigate the temporal dynamics of dosage compensated transcription in the early *Drosophila* embryo, we utilised the MS2-MS2 coat protein (MCP) system to track nascent transcription at single cell resolution in live embryos. CRISPR genome editing was used to introduce 24 copies of the MS2 loops into the large first intron of the endogenous *sog* and *mew* genes (Figure 1A). Insertion of the loops via CRISPR genome editing into the *sog* and *mew* genes did not alter expression of these genes and had little effect on viability (Figure S1). For *hnt* and *gt* we utilised previously reported fly lines with 24 MS2 loops inserted into the 5’ UTR and 3’ UTR sequences, respectively^30,31^ (Figure 1A).

**Figure 1.**
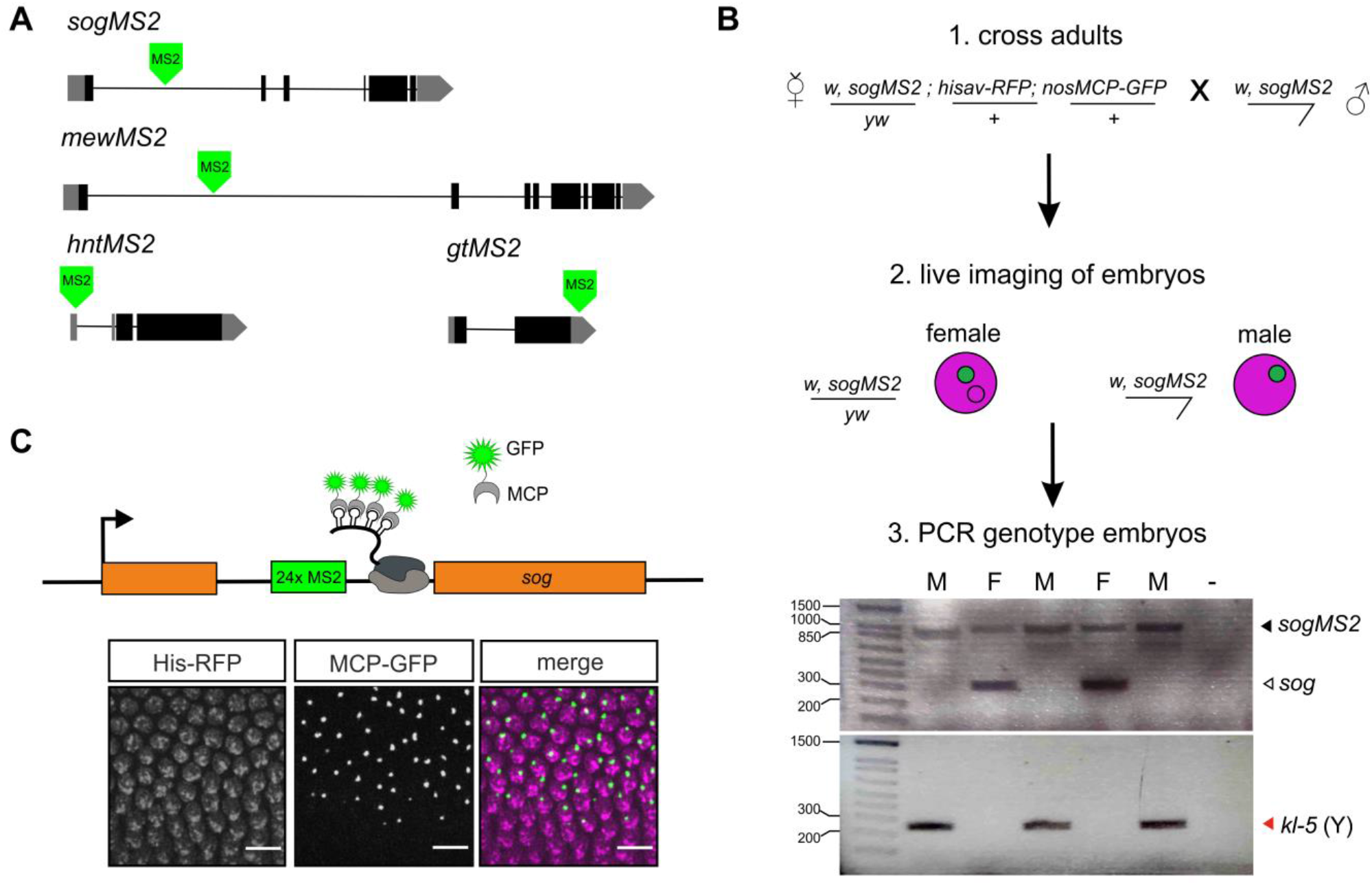
Live imaging of dosage compensated transcription in the early embryo. (A) Schematics showing the position of the 24xMS2 loops in each of the four genes used for live imaging. The *hntMS2* and *gtMS2* lines have been described previously.^30,31^ (B) Overview of the imaging and embryo sexing pipeline. The cartoon shows a female embryo with a single active TS as used for the analysis, although some female embryos imaged had 2 active TSs. The gels show representative results for PCR genotyping of individual *sogMS2* embryos to identify their sex using *sog* primers flanking the 24xMS2 loops for the X chromosome and *kl-5* primers for the Y chromosome. Female embryos have two bands for the *sog* primers as they are heterozygous for the MS2 insertion (black arrowhead) so have one unmodified *sog* locus (white arrowhead). PCR product sizes are 283bp (*sog*) and 248bp (*kl-5*), full DNA ladder sizes can be found in the Methods. (C) Top: cartoon showing that active transcription is detected by MCP-GFP binding to the MS2 loops in the mRNA as Pol II transcribes the gene. Bottom: A still from a live imaging movie corresponding to a region from the full field of view of a *sogMS2* embryo labelled with His-RFP (magenta) and the nascent transcription sites marked by MCP-GFP fluorescence (green). The border of the expression domain is visible, showing active nuclei in the presumptive neuroectoderm and inactive nuclei in the mesoderm. Scale bar is 10μm. See also Figure S1.

Females expressing MCP-GFP, His-RFP and the endogenous X chromosome gene-MS2 were crossed to males carrying the same endogenous gene-MS2 insertion (Figure 1B). Live imaging of embryos from this cross revealed bright spots of fluorescence in nuclei corresponding to nascent transcription foci of the genes of interest, where the MCP-GFP is bound to the 24xMS2 loops within the nascent mRNA transcripts (Figure 1C). Some of the embryos imaged were female embryos with 2 active MS2 transcription sites, however only movies of embryos carrying a single copy of the MS2 modified gene were analysed. After imaging each embryo was removed from the imaging dish and genomic DNA was extracted and PCR amplified using X and Y chromosome specific primers to sex the embryos (Figure 1B). For each gene the nascent transcription site (TS) fluorescent signals were visible in nuclei within the expected expression domain; the fluorescence intensity of each TS is proportional to the number of transcribing Pol II. A still from a movie from a *sogMS2* embryo is shown in Figure 1C.

### Transcriptional activities of X chromosome genes in male and female embryos

For each gene and embryo sex, nascent TSs within the expression domain were imaged live in three replicate embryos. A representative movie is shown for each gene in male and female embryos in Videos 1-8. The timing of transcription in each embryo was related to developmental time using the onset of nc14 as a reference point. Each TS was assigned to a nucleus during nc14 to reveal the spatial expression domain (Figure 2A, D, G, J) and tracked over time. For *sogMS2* embryos we analysed a consistent region by selecting nuclei within a fixed distance from the middle of the expression domain in either direction (Figure 2A). This excluded nuclei undergoing repression in the ventral region by Snail and those in the more dorsal region that have limiting activator.^32^ Heat maps of the mean fluorescence intensity traces from all nuclei analysed across the 3 biological replicates for each gender show higher signals for some male nuclei (Figure 2B). Graphs of the *sogMS2* transcriptional activity show that the mean fluorescence intensity is lower for two out of the three female embryos analysed (Figure 2C). The heat maps also reveal that nuclei have a highly synchronous onset of transcription early in nc14 in both male and female embryos (Figure 2B).

**Figure 2.**
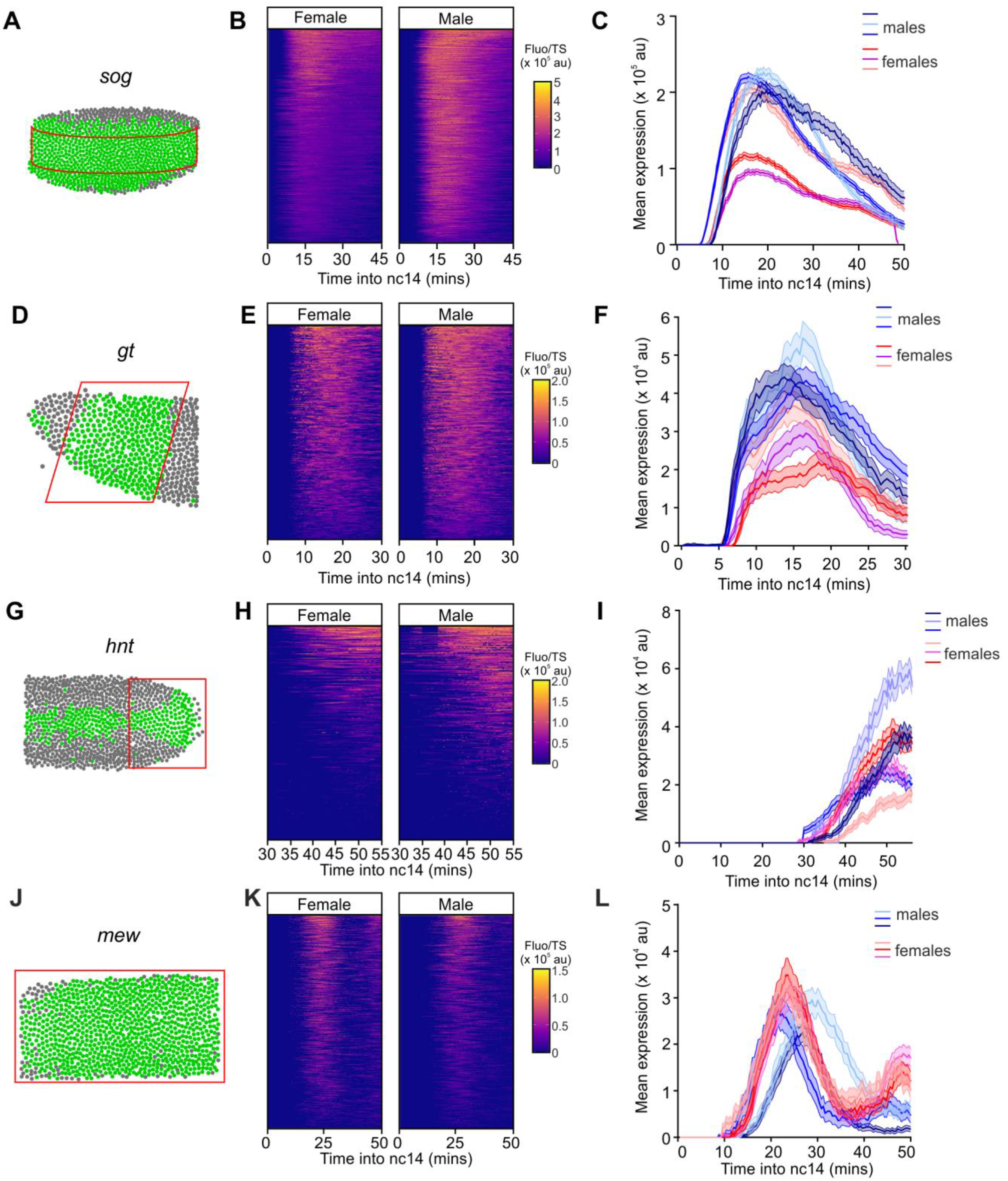
Transcriptional activities of X chromosome genes in male and female embryos. (A, D, G, J) Schematics show a representative embryo with the expression domain for the indicated gene in green, based on active nuclei from the live imaging, and the area analysed in the red box. (B, E, H, K) Heat maps show the combined individual traces for transcriptionally active nuclei from all female and male embryos during nc14. Each row shows the transcriptional activity based on mean fluorescence intensity, across developmental time in nc14. The traces are ordered by total expression. (C, F, I, L) Graphs show the mean expression based on fluorescent signals for each of the female and male embryos. Mean +/− 95% confidence intervals. See also Figure S1.

In the *gtMS2* anterior expression domain (Figure 2D), there is synchronous onset of transcription in both sexes, and a weak trend showing lower fluorescence in female embryos (Figure 2E, F). For *hntMS2*, we analysed the posterior region of the embryo (Figure 2G) where there is higher transcription and the expression domain is at its broadest. Unlike *sogMS2* and *gtMS2*, *hntMS2* transcriptional traces have low synchronicity and show a broad range of onset times in both male and female embryos (Figure 2H). Two of the male *hntMS2* embryos have similar mean TS fluorescence intensities to two of the female embryos, although the other male and female embryo have higher and lower signals, respectively (Figure 2I). For both *gt* and *hnt* some traces show fluctuating fluorescence signals, consistent with bursting (see later).

For *mewMS2* we imaged the dorsal side of the embryo and analysed all cells of the expression domain (Figure 2J). Transcription onset is stochastic and there are two peaks of transcription in both male and female embryos (Figure 2K). There is a weak trend of higher transcriptional activity in female embryos (Figure 2L). The live imaging data for the embryo replicates show some variation for each of the genes tested. This is likely biological variation, consistent with smFISH quantitation of *gt* mRNAs revealing a spread of the mean total mRNA number/cell for different embryos of the same sex (Figure S1E-G). The similar range of values between the sexes is consistent with dosage compensation occurring.^29^ Substantial fluctuations in mRNA numbers between embryos analysed as a time series has also been reported.^33^ Overall, these live imaging data suggest that there are some differences in the mean transcriptional activities between male and female embryos for the four X chromosome genes tested.

### Dosage compensated genes are not transcribed with a faster Pol II elongation rate in males

Enhanced Pol II elongation in males has been proposed to underpin dosage compensation.^18,19^ Therefore, we next estimated Pol II elongation rates from our MS2 data in male and female nc14 embryos. An autocorrelation function has been used previously to estimate Pol II elongation time from live imaging data.^34–36^ As the fluorescent signal at each transcription site is recorded at short time intervals that capture Pol II transcribing the gene, the same MS2 mRNA with bound MCP-GFP will be present at multiple time points, resulting in successive fluorescence measurements being correlated. Therefore, the autocorrelation function decays linearly with a minimum value that corresponds to the dwell time of the transcript at the TS^34–36^ (due to transcription termination in the case of *hnt*, or splicing for *sog* and *mew*).

A representative autocorrelation trace for *sog* in female embryos is shown in Figure 3A. The median dwell times for *sog*, *hnt* and *mew* in each embryo tested are shown in Figure 3B; we did not include *gt* in this analysis as the loops are located in the 3’UTR so the dwell time is extremely short. The data show that there is no significant difference between the dwell times for *hnt* in male and female embryos, whereas for *sog* and *mew* there is a small but significant increase in elongation rate (shorter dwell time) in female embryos. Based on the gene length for *hnt*, the estimated dwell times suggest elongation rates of 2.7 and 2.8 kb/min in female and male embryos, respectively. These rates are consistent with the 1.4-3.0 kb/min range of elongation rates previously measured in the *Drosophila* embryo.^37–39^ We have not converted the *sog* and *mew* dwell times to elongation rates, as we do not know how far Pol II transcribes before the nascent mRNA is spliced, given the variation in efficiency of co-transcriptional splicing in the embryo.^40^

**Figure 3.**
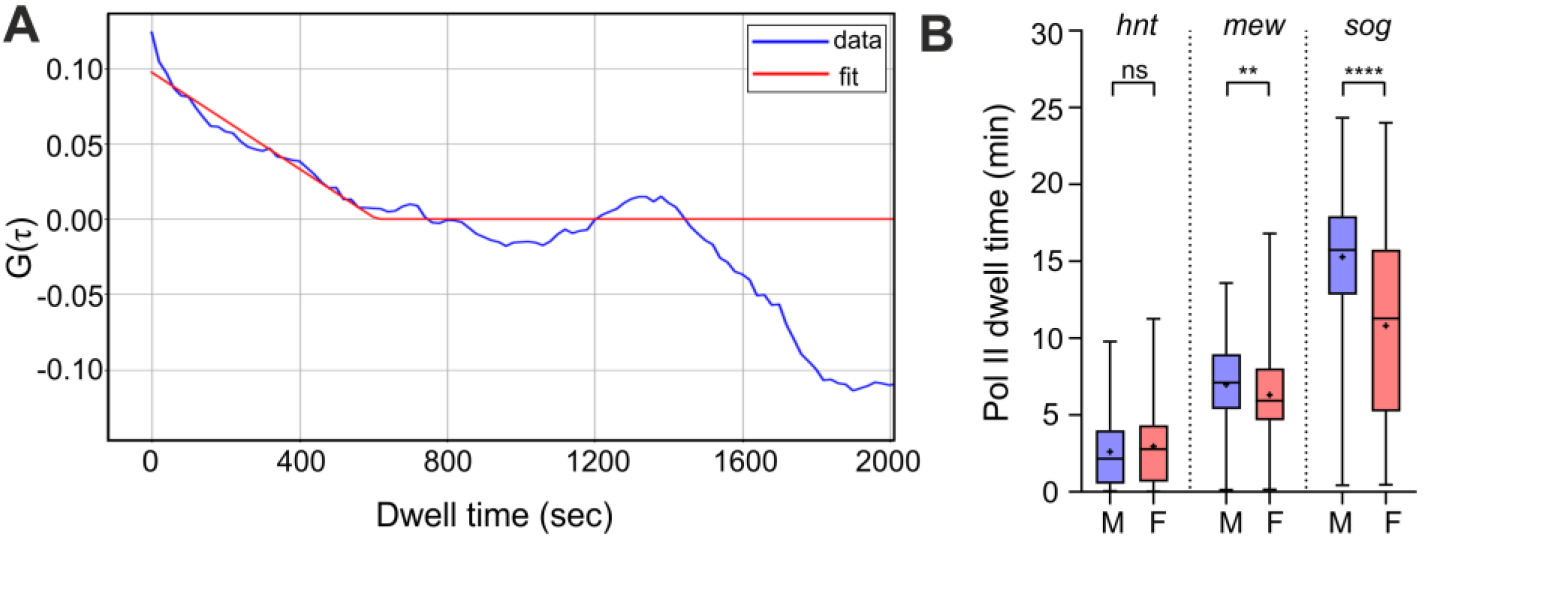
Dosage compensated genes have similar Pol II elongation rates in male and female embryos. (A) Graph shows the autocorrelation curve fit (red line) to a representative fluorescence trace (blue line) from a nucleus from a female *sogMS2* embryo. Fitting the function to the data gives the dwell time. (B) Boxplot shows the median dwell time based on the data from nuclei across replicate embryos for *hnt*, *mew* and *sog*. n= 533 (*sog* female) and n=516 (*sog* male), n=247 (*mew* female) and n=202 (*mew* male), n=194 (*hnt* female) and n=187 (*hnt* male). Boxes show 25th to 75th percentile and whiskers show range, mean is indicated with a + symbol, Welch’s t-test ** p< 0.01, **** p <0.0001, ns = not significant. See also Figure S2.

Further analysis of the dwell times from nuclei located in different regions of the *sog*, *hnt* and *mew* expression domains suggest that there is no spatial regulation of the Pol II elongation rates (Figure S2A-C). However, due to limitations with this analysis we excluded nuclei with sparse fluorescent traces (see Methods) that are typically on the edge(s) of the expression domain, so cannot rule out changes in the elongation rate in these regions. Nonetheless, as in the global analysis of Pol II elongation rate, we observe no significant difference for *hnt* between male and female embryos, whereas faster elongation rates were estimated for *gt* and *mew* in female nuclei (Figure S2A-C). Together these data do not support faster Pol II transcription in males for the dosage compensated genes tested.

### *sog* hypertranscription in males is due to a higher Pol II initiation rate

As there was no significant increase in male Pol II elongation rates, we investigated transcriptional regulation in more detail by inferring the parameters associated with transcriptional bursting. With bursty transcription, when the promoter switches from the on state to the off state fluorescence persists due to Pol II molecules transcribing the gene body. Therefore, we used a memory adjusted hidden Markov Model to infer the rates and bursting parameters from the MS2 transcriptional traces in male and female embryos.^35,41^ The model is based on a two state model of transcriptional bursting in which the promoter switches between on and off states with rates *k_on_* and *k_off_*, and initiates transcription with a rate *k_ini_*, when the promoter is in the on state. *k_on_* and *k_ini_* dictate burst frequency and amplitude, respectively, whereas burst duration is equivalent to 1/*k_off_* (Figure 4A). Promoter occupancy, based on *k_on_* and *k_off_*, is the fraction of time the promoter is in an active state^42^ (Figure 4A).

**Figure 4.**
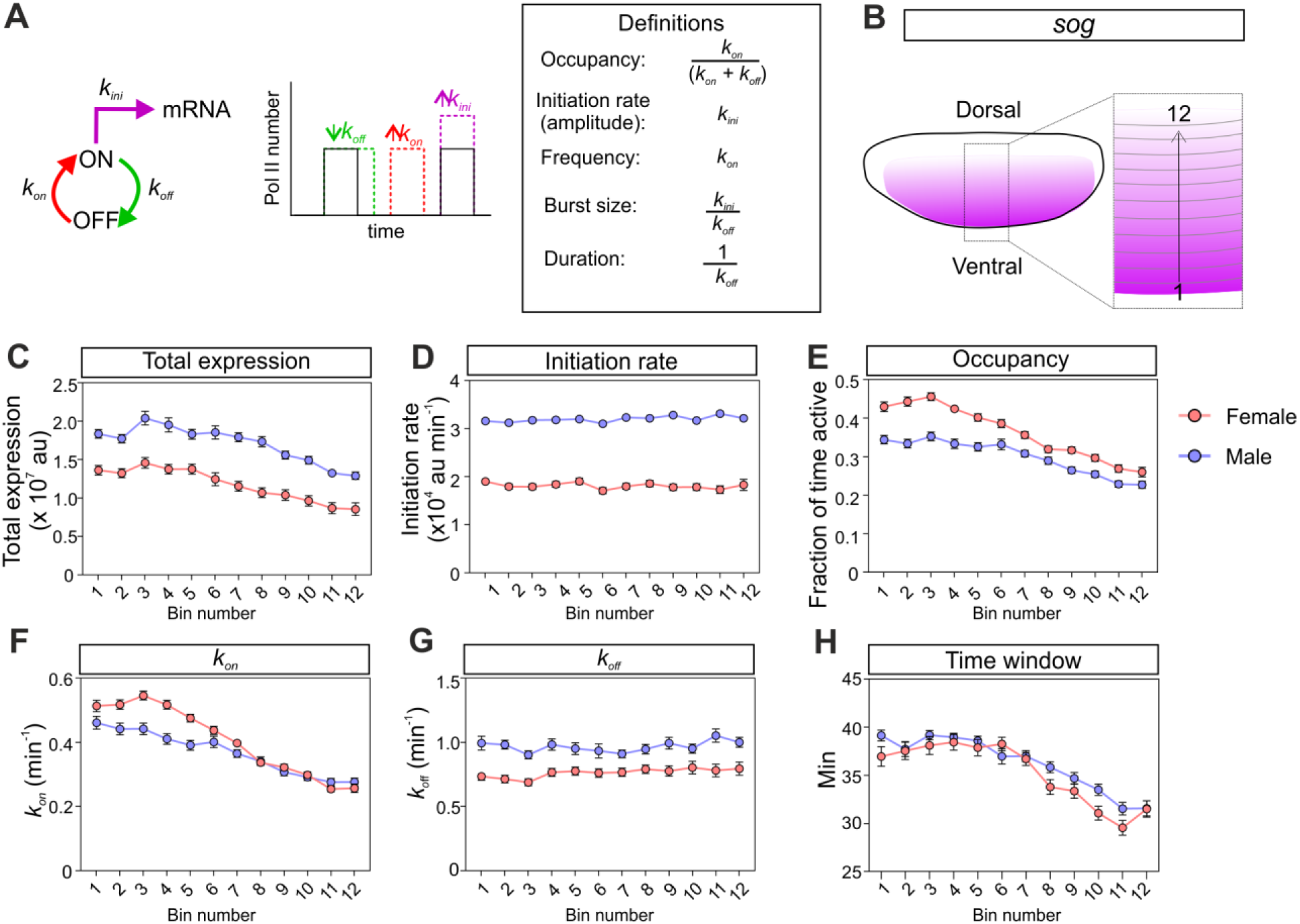
*sog* hypertranscription in males is due to a higher amplitude of transcriptional bursts. (A) Overview of the two state model of transcriptional bursting, showing *k_on_*, *k_off_* and *k_ini_* and parameter definitions. The effect of changes in these rates on bursting is shown on the graph. (B) Cartoon shows a schematic of the *sog* expression domain (ventrolateral view) with the single cell bins numbered from the ventral side of the expression domain. (C) Graph shows the mean total expression from nuclei in each single cell bin. Each male and female data point shows the data from nuclei pooled from 3 biological replicates. (D-H) Graphs show the binned single cell transcriptional parameters inferred from the *sogMS2* transcriptional traces from male (blue) and female (red) embryos: (D) Pol II initiation rate, (E) promoter occupancy, (F) *k_on_*, (G) *k_off_* and (H) transcription time window. Mean ± SEM, n= 1076 nuclei (female) and n= 1081 nuclei (male). See also Figure S3.

We used the live imaging data from nuclei in the analysed regions of the expression domain to infer burst parameters at single cell resolution in nc14 for each embryo, as described previously.^31,41^ We then pooled the data for the 3 embryos of each sex and calculated the mean parameter from nuclei divided into single-cell wide bins moving across the expression domain at nc14. For the analysis of *sog* transcription, the single cell bins are positioned along the dorsal-ventral (DV) axis and move dorsally (Figure 4B), as *sog* is activated by the Dorsal gradient.^32^ This analysis reveals that mean total expression is ~1.5 fold higher in each row of nuclei in male embryos relative to females (Figure 4C), consistent with the whole embryo trends shown in Figure 2. This magnitude of effect is in the expected range for dosage compensation which, if complete, is predicted to increase transcription 2-fold.

Pol II initiation rate is ~1.8 fold higher for *sog* in each spatial bin in male embryos (Figure 4D), whereas promoter occupancy is slightly higher in female embryos, particularly in ventral nuclei (low numbered bins) (Figure 4E). This increase in occupancy in females in ventral nuclei is due to a small increase in *k_on_* and decrease in *k_off_* (Figure 4F-G). These results suggest that the higher Pol II initiation rate on the *sog* transcription output in males is negated to some extent by the lower promoter occupancy, as the promoter spends less time in the on state in male nuclei.

The total transcription output depends on occupancy, loading rate and the transcription time window.^35,38^ We therefore calculated the time window of *sog* transcription in each nucleus by using the fluorescence data to calculate the difference between the time when transcription is first detected (t_on_) and then turns off (t_off_). This analysis reveals that male and female embryos have a similar time window in most nuclei, with only a very minor extension of the time window (up to 3 mins longer) in some of the male embryo bins (Figure 4H, S3A-B). For some nuclei, *sog* transcription is still detectable when the embryo starts to gastrulate, but the cell movements prevent continued tracking of the transcription sites. Therefore, in this analysis we have used the end of the imaging period as an estimate for t_off_ for these nuclei (see Methods). However, we were able to calculate a t_off_ value earlier than the end of the imaging time for more nuclear traces from female embryos than males (Figure S3C), suggesting that males have a longer time window for some nuclei. By multiplying Pol II initiation rate, promoter occupancy and the time window, we estimate a higher total transcription output in male embryos than females, with similar relative outputs to those observed based on the mean total fluorescent signal (Figure S3D). Together, these data suggest that dosage compensation of *sog* is mediated by a higher Pol II initiation rate in males, with the magnitude of the increase dampened by a lower promoter occupancy.

### Dosage compensated genes have elevated transcription burst amplitude in male embryos

Next, we investigated the bursting parameters for *gt*. As *gt* is activated by the Bicoid (Bcd) gradient^43^, we used spatial bins moving across the embryo from the anterior to posterior of the expression domain. Mean total expression shows a drop near the centre of the expression domain, consistent with the refinement of the broad anterior band of expression into 2 stripes.^43^ Analysis of bursting parameters reveals that there is a small (~1.4 fold) increase in total transcription in some, but not all, male nuclei that are transcribing *gtMS2* (Figure 5A). Promoter occupancy is equivalent for both sexes; although *k_on_* is higher in female nuclei in some of the bins, the higher *k_off_* in females results in a shorter burst duration (Figure 5A). Pol II initiation rate shows a small (~1.4 fold) increase across the majority of the bins in male embryos (Figure 5A). The transcription time window is longer - up to 8 mins - in many of the male nuclei bins, due to later a t_off_ (Figure S3E-G). Similar to *sogMS2*, we saw that more female than male nuclei completed transcription within the imaging time period consistent with males having an extended transcriptional window (Figure S3H). Multiplying promoter occupancy, Pol II initiation rate and the time window predicts the spatial trends of mean total expression across the expression domain in male and female *gtMS2* embryos (Figure S3I). In summary, small increases in Pol II initiation rate and, for some nuclei, the time window of transcription lead to modest increases in *gt* expression in male embryos.

**Figure 5.**
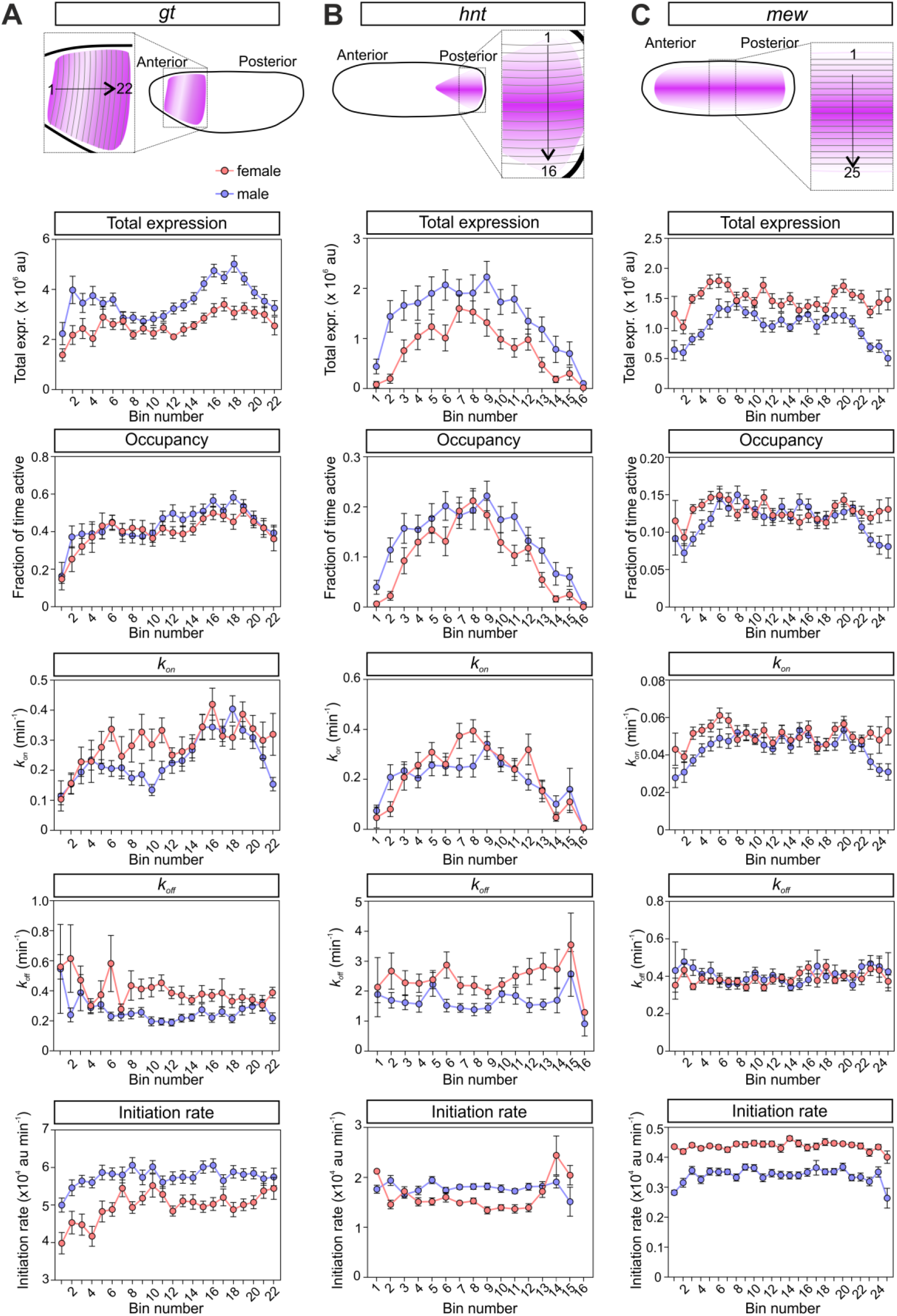
Differences in burst amplitude are associated with sex specific changes in transcription. (A) Cartoon shows the *gt* expression domain with the numbered single cell bins used in the analysis. The graphs show the mean total expression, promoter occupancy, *k_on_*, *k_off_* and Pol II initiation rate in each bin from nuclei from male (blue) and female (red) embryos. The data and single cell parameters from nuclei in 3 biological replicate embryos of each sex were pooled and reported in the indicated bins. (B, C) As in (A), but the data are shown for *hnt* (B) and *mew* (C). Mean ± SEM, n= 437 (*gt* female) and 675 nuclei (*gt* male), n= 387 (*hnt* female) and 420 (*hnt* male), n= 1067 (*mew* female) and 894 (*mew* male). See also Figure S3.

As *hnt* transcription is activated by Dpp signalling^44^, the single cell parameters were grouped in dorsal-ventral bins moving across the dorsal midline (Figure 5B). As *hnt* transcription starts late in nc14 and peak expression is only reached late in the imaging period (Figure 2I), we were unable to accurately estimate the transcription time window. Mean expression is ~1.5 fold higher in male embryos, due to small increases in both promoter occupancy (based on lower *k_off_*) and Pol II initiation rate (Figure 5B). The *hnt* parameters inferred here in the posterior of the embryo are similar to those reported previously for *hnt* transcription in nuclei in the centre of the expression domain.^31^

For *mew*, we analysed the single cell parameters in DV spatial bins moving across the dorsal midline (Figure 5C). This analysis reveals that, in contrast to the other X chromosome genes, mean expression is slightly higher in female embryos. This appears to be driven by an increased Pol II initiation rate, which is higher in female nuclei in all of the bins. In contrast, promoter occupancy and *k_on_* only show small increases at each end of the expression domain in female nuclei, whereas *k_off_* is unchanged (Figure 5C). Together, the single cell bursting parameter data suggest that *sog*, *gt* and *hnt* are dosage compensated at the transcriptional level. Promoter occupancy is modulated to different degrees for transcription of these genes, and the time window of active transcription is increased for *gt* and to a lesser extent *sog*. However, all 3 genes show increases in Pol II initiation rate in males. In contrast, Pol II initiation rate is higher in female nuclei for *mew*, which is not dosage compensated^29^, and even shows a higher transcriptional output in females.

### Burst frequency and promoter occupancy control transcriptional changes across the expression domain

Having investigated how bursting parameters change for transcription of the X chromosome genes between male and female embryos, we next used the single cell parameters to determine which parameter underpins the transcriptional changes observed spatially across each expression domain. To this end, for each embryo, we visualised mean expression and the individual parameters spatially as both heatmaps and graphs with each point representing the data from a single nucleus in the expression domain (Figure 6A, B). In addition, we calculated the correlation between mean expression and each parameter, as described previously.^31^ Analysis of the data for *sog* transcription reveals that mean expression declines in nuclei positioned more dorsally in the expression domain (Figure 6A, B), consistent with reduced levels of the Dorsal activator.^32^ Testing the correlation between mean expression and the different parameters reveals that promoter occupancy is most correlated, whereas there is little correlation between the mean expression profile across the expression domain and Pol II initiation rate, which is largely unchanged (Figure 6B, correlations for the other biological replicate embryos are shown in Figure S4A). Consistent with occupancy being highly correlated, both *k_on_* and *k_off_* are the parameters that are next most strongly correlated with mean expression (Figure 6B, Figure S4A).

**Figure 6.**
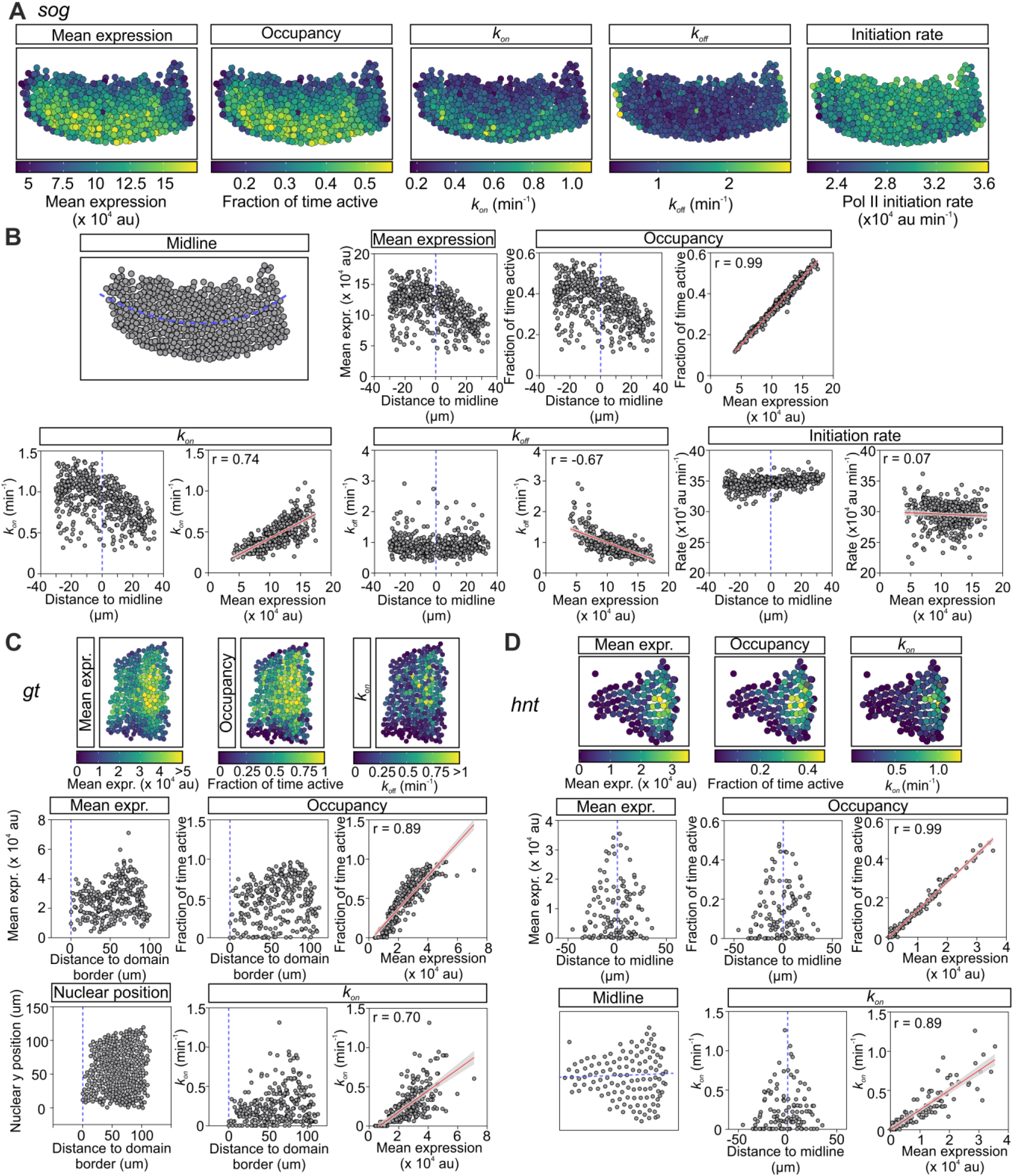
Changes in burst frequency and promoter occupancy underpin the transcriptional changes across the expression domain. (A) Heatmaps show the *sogMS2* expression domain with nuclei coloured as in the associated key for mean expression, promoter occupancy, *k_on_*, *k_off_* and Pol II initiation rate. (B) Schematic shows the *sogMS2* expression domain with its midline marked in blue. Mean expression for each nucleus in the embryo is plotted based on its position from the expression domain midline with ventral (negative) and dorsal (positive) distances. For promoter occupancy, *k_on_*, *k_off_* and Pol II initiation rate, the left graph shows the value for each nucleus plotted against expression domain position. The right graph shows the correlation between the indicated parameter and mean expression, based on the data from each individual nucleus in the expression domain for a single embryo. (C, D) As in (A, B) except the data are for *gtMS2* and *hntMS2*, and only the graphs for mean expression, promoter occupancy and *k_on_* are shown. Data for the other *gtMS2* and *hntMS2* parameters and for *mew* are shown in Figure S4. Data represent n = 475 nuclei (*sog*), 280 nuclei (*gt*) and 122 nuclei (*hnt*). See also Figure S4.

Analysis of the *gt*, *hnt* and *mew* single cell parameters across the expression domain and their correlation with mean expression also reveals that promoter occupancy is most highly correlated, to the extent that it can accurately predict the mean expression profile in each embryo (Figure 6C, D, Figure S4B-G). For all genes, *k_on_* is strongly correlated, with *k_off_* for *mew* also showing strong correlation with mean expression (Figure 6C, D, Figure S4B-G). Together, these data show how different bursting parameters are modulated to alter the transcription output in embryos in distinct ways. Our data suggest that nuclei respond to activator concentration through changes in promoter occupancy and burst frequency (*k_on_*), whereas sex specific modulation of the Pol II initiation rate further defines the transcription output.

## Discussion

Here we use live imaging to analyse the transcriptional burst kinetics for four X chromosome genes at single cell resolution. For the 3 genes previously shown to be dosage compensated^29^, we detect significantly higher mean transcriptional activity in all (*sog*) or many (*gt* and *hnt*) of the single cell bins across the expression domain in male embryos. In contrast *mew*, which was chosen as a negative control since it is not compensated^29^, shows higher transcription in female embryos. The reason for this difference in *mew* transcription in female embryos is currently unclear, but it is possible that X chromosome genes which are not compensated are under transcribed in males. The one bursting parameter in common to all these sex specific changes in transcription output is Pol II initiation rate, suggesting that control of burst amplitude underpins the transcriptional changes associated with dosage compensation. We find little change in Pol II initiation rate across the expression domain in response to changes in activator concentration, but instead it appears to be differentially tuned depending on embryo sex.

It has been shown for transcription of gap genes in *Drosophila* nc13 embryos that the initiation rate is constant for the different gap genes tested and at all positions across the expression domains.^42^ While *k_on_* and *k_off_* have been reported to be regulated in response to differing concentrations of transcription factors and cell signals^45,46^, examples where Pol II initiation rate is modulated are rarer. One example is during the refinement of the seven *even-skipped* stripes in the *Drosophila* embryo. Following the onset of their transcription, both *k_on_* and *k_ini_* are upregulated in the centre of each stripe as their expression domains narrow.^47,48^ Modulation of Pol II initiation rate also occurs at a global level to scale transcription to changes in cell size in *Schizosaccharomyces pombe*. In this model, genes compete for limiting Pol II and the amount of chromatin bound Pol II increases with cell size.^49^ Therefore, as we find different Pol II initiation rates in male and female embryos for the four X chromosome genes studied, we speculate that regulation of Pol II initiation rate may be a strategy primarily deployed by the cell to modulate transcriptional responses at a whole chromosome or transcriptome level.

Recent analysis of dosage compensation in the embryo has revealed that there is a maternal MSL subcomplex composed of MSL1, MSL3 and MOF, with the latter subunit acetylating H4K16 on all chromosomes in ovaries and pre-blastoderm embryos. This active mark is enriched at promoters prior to zygotic genome activation and increases nucleosome accessibility, priming genes for subsequent activation.^20^ The *msl-2* mRNA is detectable in both sexes of embryo at nc13 and continues to accumulate until the mRNA level declines in females at mid-nc14 but increases further in male embryos.^29^ MSL-2 protein was detected as diffuse X chromosome staining in male nc14 embryos, consistent with the canonical MSL complex becoming active at this stage. Moreover, knockdown of maternal MOF results in a reduction of transcription from all chromosomes at nc14, but there is a stronger downregulation of transcription from genes proximal to HASs in males that is not observed in female embryos.^20^ Given this timing, the canonical MSL complex could mediate the increased transcription we observe in nc14 for the dosage compensated genes. However, there is also the inverse dosage model of dosage compensation in which the MSL complex has no direct role in X chromosome transcription, based in part on considerations around normalisation and interpretation of genomic data from MSL loss-of-function studies.^15^ We note that analysis of live transcription of X chromosome and autosome genes in MSL mutants may help resolve whether or not the MSL complex acts directly on X chromosome transcription.

Non-canonical dosage compensation has also been proposed in the early *Drosophila* embryo^29,50–52^, which is Sex-lethal (Sxl) dependent but MSL independent.^50,51^ Based on the observation that multiple Sxl binding sites are more prevalent in the 3’ UTR of mRNAs transcribed from the X chromosome rather than the autosomes^52,53^, a model was suggested whereby Sxl destabilises or represses translation of mRNAs in female embryos.^52^ Additionally, miRNAs have been implicated in genomic balance and dosage compensation.^54^ It is possible that transcriptional and post-transcriptional mechanisms function together to equalize mRNA levels between males and females, as the transcriptional changes we observe (~1.5 fold) are lower than the complete dosage compensation suggested by RNA-seq.^29^ There is a precedent for this, as increased stability of mRNAs from the X chromosome compared to the autosomes has been proposed to function with hypertranscription in mammalian cells to allow dosage compensation.^55^ Recently, zygotic mRNA half-lives have been estimated in female *Drosophila* embryos.^56^ This approach could be used to determine whether dosage compensated X chromosome mRNAs are more stable in male embryos.

In terms of an initiation-^24^ vs elongation-based^18,19^ model of transcriptional hyperactivation in males, the telegraph model cannot distinguish between the changes in Pol II initiation rate being due to effects on Pol II recruitment or pause release. However, recent kinetic data from *Drosophila* and human tissue culture cells suggest that paused Pol II does not represent an essential state in between the off and permissive on states. Instead, pausing is a rare alternative off state, which cannot be captured as a distinct off state for all promoters, but is relatively long lived when it exists.^57,58^ Based on this, we favour a recruitment-based model, although further studies are required to address this. As well as studying additional X chromosome genes, models that incorporate an additional pausing state^57,58^ could be used. Our data do not support faster Pol II elongation on the bodies of the genes tested in males, as suggested previously^18,19^, with the opposite effect observed for *sog* transcription. However, as we only performed this analysis for *sog* and *hnt* as examples of compensated genes, it is possible that other male X chromosome genes are transcribed at a faster speed than in female embryos.

In addition to altered Pol II initiation rate, we also detect a small increase in the time window of active transcription for *sog* and *gt* in male embryos. This is due to a later t_off_, with transcription still active at the end of our analysis period for more male than female nuclei. Modulation of the transcriptional time window is critical for generating the *eve* stripe 2 pattern, with control of the window arising from different off times.^35^ Tethering elements have recently been described that mediate long range enhancer-promoter interactions and associations between the promoters of paralogous genes that allowing coupling of transcription dynamics.^59,60^ Loss of tethering elements alters the timing of activation and bursting dynamics.^60^ CLAMP and GAF, which both recruit the MSL complex^61^, bind tethering elements^60^, raising the possibility that CLAMP/GAF on the male X may influence the transcription time window and bursting. Alternatively, the MOF-deposited H4K16ac and action of the canonical MSL complex at nc14^20^ may facilitate nucleosome accessibility on the male X for longer.

Recently the *roX* RNAs and MSL2, which has an intrinsically disordered C terminal domain, have been found to form a stable X chromosome territory in males, which has many features of a phase separated condensate.^14^ We suggest that the male X chromosome territory concentrates Pol II, increasing the number of molecules available for transcription of X chromosome genes, thereby elevating the Pol II initiation rate and potentially the time of active transcription. X chromosome genes that are not compensated, such as *mew*, may be excluded from or have an unfavourable position, in the male X territory. However, further work is required to test this model and we note that Pol II exclusion from the inactive X chromosome during dosage compensation in mammals does not depend on biophysical compartmentalisation.^62^

We also investigated how bursting parameters are regulated in space across the expression domain. We find higher mean transcription in areas where there is increased activator concentration, e.g. in *sog* ventral nuclei or *gt* anterior nuclei.^32,43^ Our data reveal that regulation of promoter occupancy, the proportion of time the promoter is active, underpins the observed transcriptional changes across the expression domain of X chromosome genes. This parameter is also tuned to establish the transcription profiles of the *Drosophila* gap genes at nc13^42^ and nc14^30^, and the response of target genes to different BMP signalling levels.^31^ Our data suggest that occupancy is primarily modulated by changes in burst frequency (*k_on_*) in response to activator concentration. This is consistent with other reports of transcription factor concentration regulating burst frequency^45^, due to a reduction in the search time for the enhancer.^36^ *k_off_* also negatively correlates with mean expression of *sog* and *mew* in particular. Burst duration (1/*k_off_*) depends on the transcription factor dwell time^63^, suggesting that dwell time differs positionally across the expression domain, potentially due to cooperative interactions with another more localised transcription factor. Regulation of burst duration has been proposed to mediate the transcriptional response to different levels of Notch signalling.^64,65^ Overall, our data suggest that the transcription output of X chromosome genes depends on two tiers of inputs. Parameters such as burst frequency are locally tuned in nuclei within the expression domain in response to varying transcription factor inputs, whereas burst amplitude is set globally by the sex of the embryo.

### Limitations of the study

While previous genomics based studies of dosage compensation have allowed all active X chromosome genes to be studied^18,19,24^, we have focussed on transcription of only 4 X chromosome genes due to the low throughput nature of MS2 imaging. As complete dosage compensation of male X chromosome genes would result in a maximum 2-fold effect at the transcriptional level, in the context of biological variation analysis of the live imaging data is not straightforward. By binning our data spatially and pooling nuclei across biological replicates of the same sex, we present evidence that burst amplitude modulation contributes to the sex specific expression changes observed for all of the genes we studied. However, as discussed above, the two state model cannot distinguish between a higher initiation rate due to increased Pol II recruitment or enhanced pause release.

As we detected higher burst amplitude for all 3 compensated genes, we speculate that this will be a general mechanism for hypertranscription of male X chromosome genes, but further work is needed to address this. One option for inferring burst parameters for multiple X chromosome genes would be to exploit an approach for burst inference based on the two state model that was described for allele specific scRNA-seq^66^ and has recently been used with *Drosophila* scRNA-seq data.^67^ This approach can estimate *k_on_* and burst size (*k_ini_*/*k_off_*), although estimation of the individual *k_ini_* and *k_off_* parameters is less reliable.^66^ However, as noted^68^, the sparsity of reads in existing scRNA-seq data from the *Drosophila* embryo^68,69^ makes sexing the nuclei difficult. In addition, without allele specific scRNA-seq data the model requires an additional parameter, the frequency with which the bursting from the two alleles in female nuclei is coordinated. As the frequency of co-bursting is not trivial to estimate and likely changes across expression domains, inferring allele-specific burst parameters for multiple X chromosome genes from scRNA-seq data is currently challenging.

## Acknowledgements

We thank Caroline Hoppe, Jennifer Love and Sophie Frampton for comments on the manuscript, Takashi Fukaya for the *gtMS2* flies, Catherine Sutcliffe for technical help, the Bloomington Drosophila Stock Center for flies, Cambridge Fly Facility for microinjections and the University of Manchester Bioimaging and Fly facilities for support. This project was supported by a Wellcome Trust Investigator award to H.L.A. and M.R. (204832/Z/16/Z, 204832/B/16/Z).

## Author contributions

Conceptualization, L.F.B., M.R. and H.L.A.; Investigation, L.F.B, H.Z.; Writing, L.F.B. and H.L.A.; Supervision, M.R. and H.L.A.; Funding Acquisition, M.R. and H.L.A.

## Methods

### RESOURCE AVAILABILITY

#### Lead Contact

Further information and requests for resources and reagents should be directed to and will be fulfilled by the Lead Contact, Hilary L. Ashe.

#### Materials Availability

Plasmids and fly lines generated in this study are available from the Lead Contact on request.

#### Data and Code Availability

All data reported in this paper will be shared by the lead contact upon request. This paper does not report original code. The analysis code used to connect statistics files from Imaris are available on GitHub (https://github.com/TMinchington/sass, RRID:SCR_018797). The modelling software for inferring burst parameters can be found on GitHub (https://github.com/ManchesterBioinference/burstInfer). Any additional information required to reanalyse the data reported in this paper is available from the lead contact upon request.

### EXPERIMENTAL MODEL AND SUBJECT DETAILS

#### Experimental animals and crosses

All stocks were grown and maintained at 20°C and raised at 25°C for experiments on standard fly food media (yeast 50g/L, glucose 78g/L, maize 72g/L, agar 8g/L, 10% nipagen in EtOH 27ml/L and propionic acid 3ml/L).

The following fly lines were used in this study, y^1^ w* (BDSC Stock #6599, RRID:BDSC_6599), y^1^ w*; P{His2Av-mRFP1}II.2; P{nos-MCP.EGFP}2 (BDSC Stock #60340, RRID:BDSC_60340), *y*^1^ *w*^1118^ M{vas-Cas9}ZH-2A; 24xMS2-*hnt*,^31^ *gtMS2*,^30^ *w*^1118^ 24xMS2-*sog* (this study), *w*^1118^ 24xMS2-*mew* (this study), *w*^1118^; PBac{vas-Cas9}VK00027 (BDSC Stock #51324, RRID:BDSC_51324), y^1^ w^67c23^; MKRS, P{ry[+t7.2]=hsFLP86E/TM6B, P{w[+mC]=Crew}DH2, Tb^1^ (BDSC Stock #1501, RRID:BDSC_1501).

For all live imaging experiments His2Av-mRFP; nos-MCP-EGFP virgin females were crossed to males carrying the target gene-MS2 locus. F1 virgin females of genotype gene-MS2/+ His2Av-mRFP/+ nos-MCP-EGFP/+ were crossed to gene-MS2 males to obtain F2 male and female embryos (Figure 1B) containing the gene-MS2 locus and maternally loaded His2Av-RFP and MCP-EGFP. The male and female F2 embryos analysed have one copy of the gene-MS2 insertion.

### METHOD DETAILS

#### CRISPR of 24X MS2 loops into endogenous loci

24xMS2 loops^70^ (from pCR4-24XMS2SL-stable, RRID: Addgene_31865) were inserted into the first intron of *sog* and *mew* using two guide RNAs and one-step CRISPR Cas9 genome engineering^71–73^. Briefly, two guide regions were chosen within the first intron of the *sog* and *mew* genomic loci at a central position to avoid splice sites. A double stranded donor plasmid was constructed containing the intronic region that was removed between the two guides, the 24xMS2 loop cassette and a DsRed marker (from pHD-DsRed, RRID:Addgene_51434) inserted using a ClaI site for *mew* and AccII site for *sog*. The PAM sequences within each donor plasmid were mutated using site directed mutagenesis with Pfu Turbo (Agilent, Cat# 600250) to avoid targeting of the donor plasmid by Cas9 nuclease. Both the donor plasmid and the two guide RNA plasmids (pU6-BbsI-chiRNA, RRID:Addgene_45946) for each gene were injected into Cas9 embryos (BDSC Stock #51324, RRID:BDSC_51324) by the Cambridge Fly Facility. Oligonucleotide sequences for guide RNAs are listed in Table S1. Successful transformants were selected using the DsRed marker, which was subsequently removed by crossing to a Cre recombinase stock (BDSC Stock #1501, RRID:BDSC_1501) and screening for loss of the marker in the next generation. All primer sequences are listed in Table S1.

#### Single molecule fluorescent in situ hybridisation

2-4 hour embryos were fixed as previously described^74^ and stored in methanol at −20°C until required. Fixed embryos were placed in Wheaton vials (Sigma, Cat# Z115053-12EA) for FISH as described previously^31^. Embryos were probed for the mRNA target using smiFISH fluorescent probes designed to exonic sequences of *gt*, *sog* and *mew* with X or Z flap sequences^75^ and secondary detection probes labelled with Quasar 570 or 670 fluorophore (all probe sequences are listed in Table S1). Mouse α-Spectrin antibody (DSHB, 3A9 (323 or M10-2), RRID:AB_528473) incubation overnight at 4°C was used with a secondary Goat anti-Mouse IgG (H+L) Cross-Adsorbed Secondary Antibody, Alexa Fluor 488 (Thermo Fisher Scientific, Cat# A-11001, RRID:AB_2534069) for 2 hours at room temperature to stain the membrane. DAPI (New England Biolabs, Cat# 4083) was added to the embryos in the second of the final four washes of the protocol at a concentration of 1:1000 and embryos were mounted onto slides in Prolong Diamond (Thermo Fisher Scientific, Cat# P36961) to set overnight before imaging.

#### PCR assay to identify embryo sex

After live-imaging, individual embryos were carefully picked off the imaging dish and stored at −20°C in individual tubes. DNA was extracted from single embryos by crushing them in 50μl of squishing buffer (10mM Tris-Cl pH8.2, 1mM EDTA, 25mM NaCl, 200ug/ml Proteinase K) and incubating at 25°C for 25 mins followed by 2 min incubation at 95°C to inactivate the Proteinase K^76^. PCR amplification of DNA was performed using GoTaq (Promega, Cat# M7123) or Phusion (New England Biolabs, Cat# M0530) DNA polymerase following the manufacturer’s protocol. Primers were used that flanked the 24XMS2 cassette insertion and amplified the *kl-5* gene on the Y chromosome (primer sequences are listed in Table S1), which allowed detection of the presence of the MS2 loops on the X chromosome and either an unmodified locus (female embryos) or the *kl-5* gene (male embryos). Due to the repetitive nature of the MS2 loops, the primers that flank the MS2 insertion produce a PCR product that can vary from ~1-1.6kb therefore the PCR product from the unmodified locus in combination with the male Y chromosome band was used primarily to identify embryo sex. PCR reactions were performed in triplicate for each embryo. The 1 Kb Plus DNA Ladder (Thermo Fisher Scientific, Cat# 10787018) was used in Figure 1 with the following band sizes shown on the gel (100, 200, 300, 400, 500, 650, 850, 1000, 1500bp).

#### Viability assay

*y w / y^+^ w sogMS2* females were crossed to *y^+^ w sogMS2 / Y* males. Each replicate experiment consisted of six vials, each containing thirty larvae. The survival of adult males was measured by calculating the relative proportion of *y* and *y^+^* males emerging. The same crossing scheme was used to assess *mewMS2* viability.

#### Confocal microscopy of fixed embryos

An Andor Dragonfly200 spinning disk upright confocal microscope with a 40x / 1.30 HCL pL Apochromat objective was used to acquire smFISH images of *sog* and *mew* in fixed embryos. Samples were excited using 405nm (10%), 488nm (11%) and 637nm (10%) diode lasers respectively. Images were collected with an iXon EMCCD camera (1024 × 1024) with a gain of 180 for 130ms of multiple Z stacks at system optimised spacing.

For experiments quantifying *gt* mRNA counts (Figure S1E) a Leica TCS SP8 gSTED confocal was used using a 100x/ 1.3 HC PI Apo Cs2 objective at 0.75X zoom. Confocal settings were 1 airy unit pinhole, 400 Hz scan speed with bidirectional line scanning and a format of 4096 x 4096 pixels. Laser detection settings were collected as follows: PMT detector DAPI excitation at 405nm (7%, collection: 415-470nm); Hybrid SMD Detectors: AlexaFluor 488 excitation at 490nm (12%, collection: 500-540nm), Quasar 570 excitation at 548nm (20%, collection: 558-640nm) with 1-6ns gating. All images were collected sequentially and optical stacks were acquired at system optimised spacing. Imaging of the membrane stained with anti-Spectrin at the mid-sagittal plane of the embryo with 40x objective at 0.75X zoom and 1024 × 1024 format was used to measure the average length of membrane invagination from at least 5 cells. These measurements were used to select embryos of a similar age in early nuclear cycle 14 (~5μm membrane invagination). For all analysis, 6 separate embryos of each sex were imaged and quantified as independent replicates.

#### Live imaging Microscopy

Embryos were laid on apple juice agar plates for approximately 1 hour and embryos were collected and dechorionated in 50% bleach solution (2.5% final concentration of sodium hypochlorite solution diluted in water). Preparation of embryos for live imaging was performed as described^77^, with embryos mounted onto a heptane glue coated coverslip (Scientific Laboratory Supplies, Cat# MIC3110) and inverted over a coverslip bridge in a 7:1 ratio mix of 700:27 halocarbon oil (Sigma, Cat# H8773 and Cat# H8898) on the membrane of a Lumox dish (Sarstedt AG & Co, Cat# 94.6077.305). Images were collected on an Andor Dragonfly200 spinning disk upright confocal microscope with a 40x / 1.30 HCL pL Apochromat objective. Samples were excited using 488nm (11%; *sogMS2*, *gtMS2* and *mewMS2* or 13%; *hntMS2*) and 561nm (6%) diode lasers via Leica GFP and RFP filters respectively. Images were collected simultaneously using dual camera imaging with Zyla 4.2 Plus sCMOS (2048 × 2048) and iXon EMCCD camera (1024 × 1024) with a gain of 180 and binning [2X and 1X respectively] for 130ms. For each movie a total of 50 Z stacks at 0.7μm spacing were collected using the fastest setting yielding a total Z size of 35μm at a time resolution of between 20-25 seconds on average.

#### Image deconvolution

Images were deconvolved using either the inbuilt Andor deconvolution software for the live embryo movies or Huygens professional deconvolution software by SVI (Scientific Volume Imaging, RRID:SCR_014237) for smFISH images. smFISH images of whole embryos were tiled using the Grid/Collection stitching plugin in FIJI (ImageJ) (NIH, RRID: SCR 002285).

### QUANTIFICATION AND STATISTICAL ANALYSIS

#### Live and fixed embryo image analysis

Imaris software ≥9.2.1 (Bitplane, RRID:SCR_007370) was used for nuclear segmentation and spot detection of transcription sites (TSs) in live imaging movies. Nuclear segmentation was performed using the “surface” function with tracking autoregressive motion and maximum frame gap of 5 and travel distance of 5μm. The “spots” function was used to detect TSs in 3D with a set size of 1.5μm in X/Y diameter and 5μm (*gtMS2)* or 10μm (*sogMS2, mewMS2* and *hntMS2)* in the Z direction. Multiple background spots of the same size as the TS were added manually to every third time point to be used for background correction of the fluorescent signal. For fixed embryos the same nuclear segmentation (without tracking) and spot detection was used with spots of size 0.2 μm used to detect single mRNAs instead of TSs. All statistics were exported and the custom sass python script assigned the TS spots to nuclei across time with background correction or mRNA spots to nuclei at a single time point (Github; https://github.com/TMinchington/sass). Further statistical and data analysis was performed in R (version 4.1.2), Python and GraphPad Prism (9.1.2, RRID: SCR 002798). For all instances where nuclei were binned, bins used were 5μm in width (approximately one nucleus in width).

#### Autocorrelation estimation of elongation time

To determine the rate of Pol II elongation we used the data from the MS2 movies and determined the autocorrelation function of fluorescent traces^36^ known as G(*τ*)

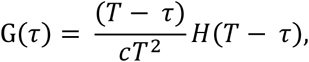

where *T* is the dwell time, *τ* is the autocorrelation delay, *c* is the initiation rate of Pol II and *H* is the Heaviside step function. This function calculates the degree to which a fluorescence signal at one time point *F(t)* is correlated to a lagged signal of itself *F(t -τ*) as a function of *τ*. Therefore the fluorescence signal at any given time point *t* will be correlated with an earlier fluorescence value *F(t - τ)* when *τ* < *T*. Under this condition, the two time points will have shared Pol II on the gene and will therefore be correlated. As *τ* increases, the correlation from the shared Pol II on the gene will decline linearly until it reaches a transition point at which point it will be equivalent to *T* and is taken as the time for Pol II to traverse the gene or for splicing to occur. The dwell time was calculated for fluorescence traces individually and the median value for each embryo calculated. For this analysis we used transcriptional traces for each gene with moderate to high transcriptional activity, due to difficulties with the analysis for the sparse traces from low expressing nuclei. The elongation rate for *hnt* was calculated by dividing by the gene length with the dwell time.

#### Modelling transcriptional parameters

MS2 fluorescence traces from all nuclei were used to infer promoter states using a memory-adjusted hidden Markov model (mHMM) implemented in python with a truncated state-space approximation^41^. The model was trained on each embryo separately to generate the transcriptional parameters by sex. The global parameters obtained were the rate of promoter switching on (*k_on_*) and off (*k_off_*), the Pol II initiation rate (*k_ini_*) and promoter mean occupancy <*n*> as defined previously^42^.

Single cell parameters were determined from the mHMM for each embryo^41^. Single cell parameters were combined for all female and male replicates and plotted into 5μm bins across the expression domain for each gene (Figures 4 and 5). For each replicate the single cell parameters were plotted against either the corresponding distance to the expression domain midline/border or mean expression to determine correlations (Figure 6). To determine t_on_ for a given nucleus, the first time a nucleus increases from zero was taken and for t_off_ a nucleus must be reduced to zero for 5 consecutive time points at the end of the trace. If a nucleus did not have a t_off_ due to transcription still being active at the end of the imaging period, then the final imaging time point was taken as t_off_. All statistical analysis was carried out in R (version 4.1.2), Python and GraphPad Prism (9.1.2, RRID: SCR 002798).

#### Supplemental Videos and Table

Videos S1, S2. Maximum intensity projection of a lateral view of a representative male (Video S1) and female (Video S2) embryo expressing *sogMS2* (green) and His-RFP (magenta) expression imaged with a 40x objective and 20 sec time resolution during nc14.

Videos S3, S4. As in Videos S1 and S2, except the male (Video S3) and female (Video S4) embryos are expressing *gtMS2* (green) and His-RFP (magenta).

Videos S5, S6. As in Videos S1 and S2, except the male (Video S5) and female (Video S6) embryos are expressing *mewMS2* (green) and His-RFP (magenta) and imaged as dorsal views.

Videos S7, S8. As in Videos S1 and S2, except the male (Video S7) and female (Video S8) embryos are expressing *hntMS2* (green) and His-RFP (magenta) and imaged as dorsal views.

Table S1. smFISH probe and primer sequences.

