## Supplementary Figures for "Modulation of transcription burst amplitude underpins dosage compensation in the *Drosophila* embryo"

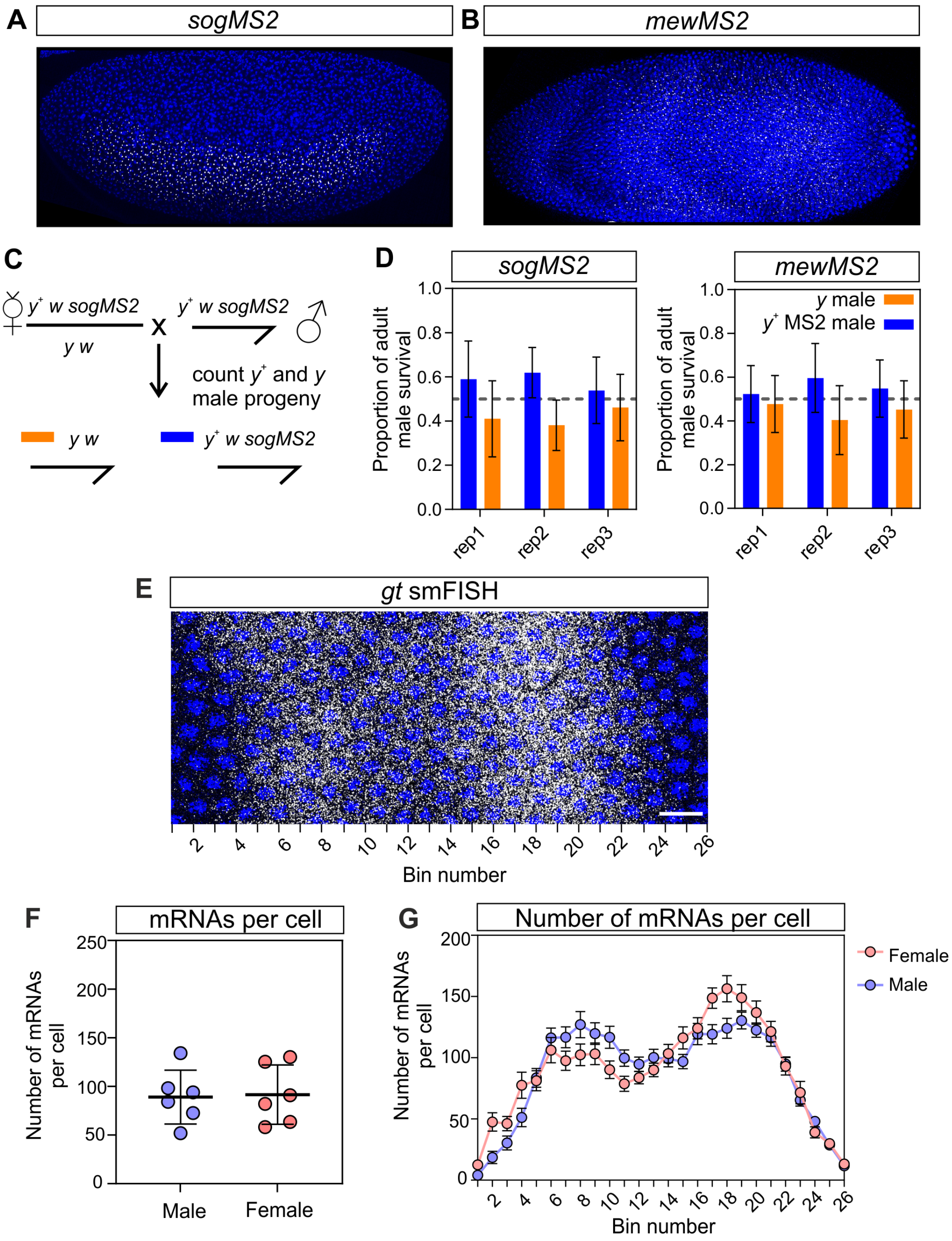


**Figure S1. Insertion of MS2 loops into the *sog* and *mew* endogenous loci has no effect on expression or viability. Related to Figures 1 and 2.**

(A) smFISH of a *sogMS2* embryo with *sog* probes (grey) and DAPI staining (blue) showing the expected expression domain. A lateral view of the embryo is shown at nc14.

(B) As in (A) except a *mewMS2* embryo is stained with *mew* smFISH probes and a dorsal view is shown.

(C) Overview of the crossing scheme to determine the viability of adult flies containing the *sog* or *mew* locus with a 24xMS2 insertion (*sogMS2* is shown as the example).

(D) Graphs show the viability of *sogMS2* or *mewMS2* males relative to those carrying an unedited X chromosome. The dotted line on the graph marks the expected proportion if viability is unaffected by insertion of the MS2 loops. Differences are not significant between control and MS2 male survival. $p > 0.05$, n= 5-6 independent experiments for each of 3 biological repeats, Holm-Sidak's multiple t-test. Mean ± SD of n = 87, 86 and 98 (*sog*MS2) and n = 68, 84 and 63 (*mew*MS2).

(E) Representative smFISH image showing *gt* mRNAs in part of a nc14 embryo, with the expression domain divided into different bins.

(F) Graph shows the mean number of *gt* mRNAs/cell for male or female embryos. Mean ± SD of n = 6 embryos of each sex.

(G) Graph shows the mean number of mRNAs per cell binned spatially across the expression domain at early nc14, bin numbers are as in (E). Mean ± SEM of all cells within the binned region. n= 6 embryos of each sex.

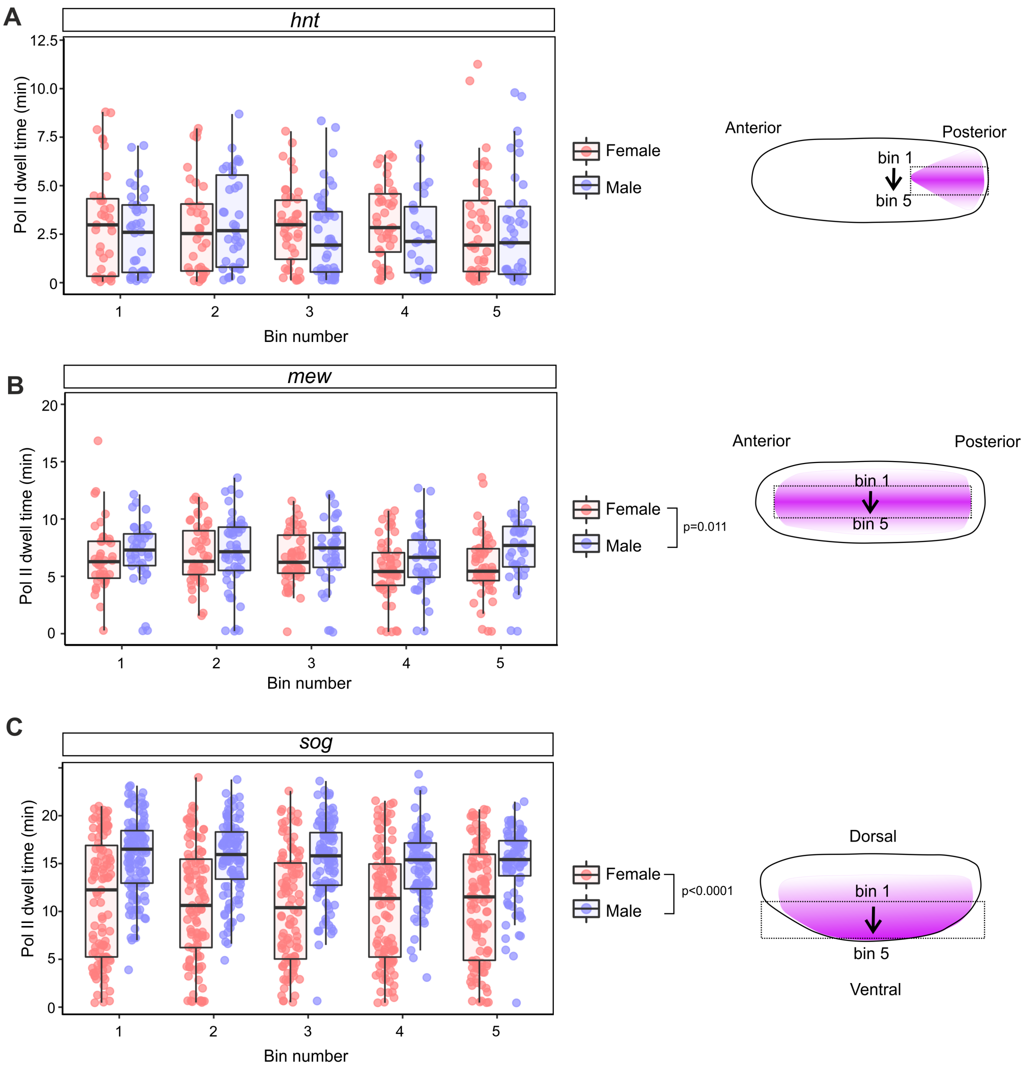


**Figure S2. Spatial analysis of Pol II dwell times in embryos. Related to Figure 3.**

(A-C) Boxplots of the Pol II dwell time per transcription site plotted for each bin across a region of the expression domain for (A) *hnt*, (B) *mew* and (C) *sog* in male and female embryos. Boxes show 25th to 75th percentile, line shows median and whiskers show 1.5x interquartile range. Bin numbers correspond to those in the cartoons. A two way ANOVA was used to determine statistical significance. Testing the effect of sex and bin number on Pol II dwell time found that males had a significantly longer Pol II dwell time for *sog* $F_{1,\ 1039} = 194.6$, $p < 0.0001$ and *mew* $F_{1,\ 439} = 6.5$, $p = 0.0114$ but not for *hnt.* There was no significant effect of the bin number on Pol II dwell time for any of the genes, consistent with no spatial regulation of elongation rate. n=194 (*hnt* female) and n=187 (*hnt* male), n=247 (*mew* female) and n=202 (*mew* male), n= 533 (*sog* female) and n=516 (*sog* male).

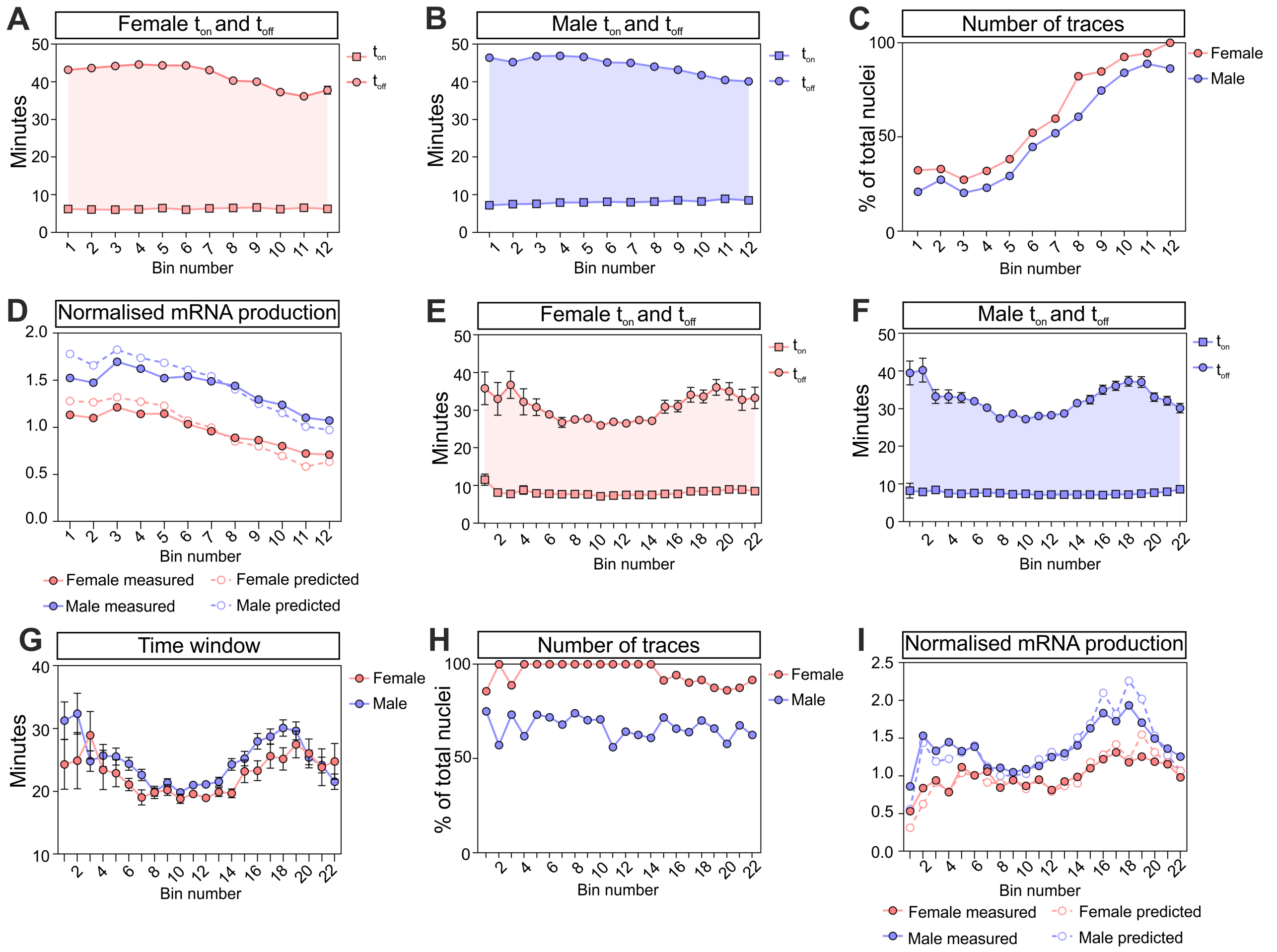


**Figure S3. The *sog* and *gt* transcription time windows in male and female embryos. Related to Figures 4 and 5.**

(A, B) Graphs show the mean values for $t_{on}$ (squares) and $t_{off}$ (circles) in nuclei from *sogMS2* female (A) and male (B) embryos. The single cell bins are as described in Figure 4B. The shaded area represents the total time window ($t_{off}$ - $t_{on}$).

(C) Percentage of *sogMS2* nuclear traces that have a $t_{off}$ within the imaging period.

(D) Graph shows the predicted total *sogMS2* expression in male and female embryos, based on multiplying Pol II initiation rate, the active time window and promoter occupancy (open circles, dashed lines) with the expression data from Figure 4C plotted for comparison (closed circles). Data have been normalised to the mean value from the female data.

(E, F). As in (A, B), except the data are for *gtMS2*. The single cell bins are as described in Figure 5A.

(G) Graph shows the mean time window estimated for *gtMS2* across single cell bins in male and female embryos.

(H) As in (C), except the data are for *gtMS2*.

(I) As in (E), except the data are for *gtMS2.* Mean ± SEM, n= 1076 (*sog* female) and 1081 (*sog* male), n= 437 (*gt* female) and 675 nuclei (*gt* male).

**A**

| Replicate | $k_{on}$ | $k_{off}$ | Occupancy | Initiation rate |
|---|---|---|---|---|
| *sog* female 1 | 0.55 | -0.75 | 0.91 | 0.17 |
| *sog* female 2 | 0.84 | -0.67 | 0.99 | 0.05 |
| *sog* female 3 | 0.70 | -0.71 | 0.98 | 0.18 |
| *sog* male 1^ | 0.74 | -0.67 | 0.99 | 0.07 |
| *sog* male 2 | 0.81 | -0.77 | 0.97 | 0.07 |
| *sog* male 3 | 0.68 | -0.63 | 0.99 | -0.08 |

**B**

| Replicate | $k_{on}$ | $k_{off}$ | Occupancy | Initiation rate |
|---|---|---|---|---|
| *gt* female 1 | 0.64 | -0.34 | 0.88 | 0.38 |
| *gt* female 2 | 0.60 | -0.43 | 0.93 | 0.20 |
| *gt* female 3 | 0.63 | -0.42 | 0.90 | 0.59 |
| *gt* male 1^ | 0.70 | -0.48 | 0.89 | 0.66 |
| *gt* male 2 | 0.74 | -0.32 | 0.87 | 0.57 |
| *gt* male 3 | 0.66 | -0.34 | 0.89 | 0.66 |

**C** *gt*

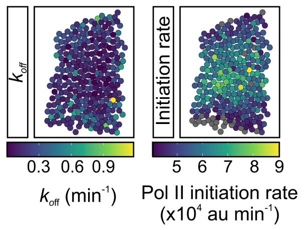


**D**

| Replicate | $k_{on}$ | $k_{off}$ | Occupancy | Initiation rate |
|---|---|---|---|---|
| *hnt* female 1 | 0.89 | -0.32 | 0.99 | 0.15 |
| *hnt* female 2 | 0.88 | -0.46 | 0.99 | 0.05 |
| *hnt* female 3 | 0.85 | -0.29 | 0.99 | 0.14 |
| *hnt* male 1^ | 0.89 | -0.21 | 0.99 | -0.11 |
| *hnt* male 2 | 0.87 | -0.28 | 0.99 | -0.12 |
| *hnt* male 3 | 0.82 | -0.53 | 0.99 | 0.04 |

**E** *hnt*

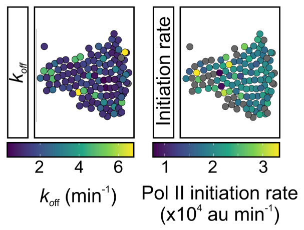


**F** *mew*

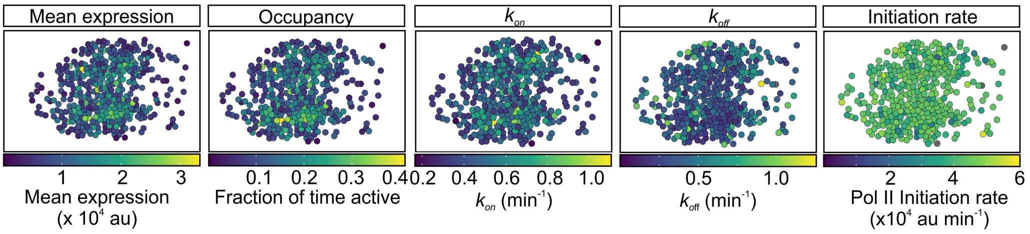


**G**

| Replicate | $k_{on}$ | $k_{off}$ | Occupancy | Initiation rate |
|---|---|---|---|---|
| *mew* female 1 | 0.65 | -0.39 | 0.97 | 0.18 |
| *mew* female 2 | 0.50 | -0.56 | 0.97 | 0.37 |
| *mew* female 3 | 0.64 | -0.51 | 0.95 | 0.34 |
| *mew* male 1^ | 0.62 | -0.48 | 0.96 | 0.12 |
| *mew* male 2 | 0.66 | -0.55 | 0.97 | 0.39 |
| *mew* male 3 | 0.48 | -0.55 | 0.97 | 0.31 |

**Figure S4. Correlations between mean expression and single cell burst parameters for all nuclei in the expression domain. Related to Figure 6.**

(A, B, D, G) Table shows the Pearson correlation coefficients between mean expression (arbitrary units) and the indicated burst parameter for all nuclei in each individual male and female embryo analysed. The correlations are shown for *sog* (A), *gt* (B), *hnt* (D) and *mew* (G). The ^ label denotes the embryo used to show the data as spatial heatmaps in panels S4C, E and F and Figure 6A, C, D.

(C, E) Spatial heatmaps from a representative *gt* (C) and *hnt* (E) embryo showing $k_{off}$ and Pol II initiation rate with nuclei coloured as in the associated key.

(F) Heatmaps from a representative *mew* embryo, with nuclei in the expression domain coloured depending on their value for either mean expression or the indicated burst parameter. The colour keys are shown below each heatmap.
