## Supplementary Table 1 for "Modulation of transcription burst amplitude underpins dosage compensation in the *Drosophila* embryo"

| **Primer/probe name** | **Sequence 5' to 3'** |
| --- | --- |
| mewint_HA_L3F | ctgtttgcaccgcgagaatt |
| mewint_HA_L3BamR | atcgGGATCCgaaagaaggttgccgctctc |
| mewint_HA_R3BglF | tacgAGATCTgctgcaaacggggtcacc |
| mewint_HA_R3R | ttggagctgaagttgttgcc |
| mewint_HA1seqF | acgattttggctggcagttt |
| mewint_HA1seqR | gaaaggtggcagggaatgtg |
| mewint_guide6Lmut_F | cgtatcgagcctttcgcttagttagcatctaag |
| mewint_guide6Lmut_R | cttagatgctaactaagcgaaaggctcgatacg |
| mewint_guide5Rmut_F | ggcatttcgagacgagtcgtttggcttcgcgc |
| mewint_guide5Rmut_R | gcgcgaagccaaacgactcgtctcgaaatgcc |
| mew_G5_S | CTTCGCTCTTGGCATTTCGAGACG |
| mew_G5_AS | AAACCGTCTCGAAATGCCAAGAGC |
| mew_G6_S | CTTCGAAGATCGTATCGAGCCTTT |
| mew_G6_AS | AAACAAAGGCTCGATACGATCTTC |
| dsred_ClaI_F | cggaatcgatccgcggacatatgcaCAC |
| dsred_ClaI_R | ctagatcgatgatctttactagtGCTCTTCt |
| sogint_HA_L1F | tagtcgttatatggccgcga |
| sogint_HA_L1BamR | atcgGGATCCacgtgtttcgtggccgacc |
| sogint_HA_R1BglF | tacgAGATCTtaagaaagagcggtccaggg |
| sogint_HA_R1R | tcttgttgctgcgttgttga |
| sogint_seqF3 | atgtggtggatggtggatgt |
| sogint_seqR3 | tacgtccgtctctttctcgg |
| sogint_HA_L1F | atcgCCATGGtagtcgttatatggccgcga |
| sogint_HA_R1R | atgcGTCGACtcttgttgctgcgttgttga |
| sogint_guide4Lmut_F | Cattggactttaagacagggtagctgcgc |
| sogint_guide4Lmut_R | gcgcagctaccctgtcttaaagtccaatg |
| sogint_guide3Rmut_F | Gatggcatcaatacctcttataccc |
| sogint_guide3Rmut_R | gggtataagaggtattgatgccatc |
| sogint_HA_L1F | atcgCCATGGtagtcgttatatggccgcga |
| sogint_HA_R1R | atgcGTCGACtcttgttgctgcgttgttga |
| sog_G4_S | CTTCGATGTATGTGCGCAGCTACCC |
| sog_G4_AS | AAACGGGTAGCTGCGCACATACATC |
| sog_G3_S | CTTCGATCGGGATCGGGTATAAGA |
| sog_G3_AS | AAACTCTTATACCCGATCCCGATC |
| dsred_AccIII_F | cggaTCCGGAccgcggacatatgcaCAC |
| dsred_AccIII_R | ctagTCCGGAgatctttactagtGCTCTTCt |
| sogMF_F | CATATCATGATCAGCGGCGG |
| sogMF_R | CCCTGGACCGCTCTTTCTTA |
| gtMF_F | TAACTGGTGGGATTGCGAGA |
| gtMF_R | TCAAGGAGGATGAGATCGCC |
| hntMF_F | ACCAAGCCTAAAACAGTGCG |
| hntMF_R | CCATGTTTTCTGCGCTCGAT |
| mewMF_F | cgagagcggcaaccttctt |
| mewMF_R | gctagcaatcaaggcaggaaa |
| kl5 F | GCTGCCGAGCGACAGAAAATAATGACT |
| kl5 R | CAACGATCTGTGAGTGGCGTGATTACA |
| mew_ex1 | CCAGCTTCTAGCATCCATGCCCTATAAGAACACCATGGTGTCATCGTC |
| mew_ex2 | CCAGCTTCTAGCATCCATGCCCTATAAGCTGAGTCACCCAAATGGATC |
| mew_ex3 | CCAGCTTCTAGCATCCATGCCCTATAAGCACTGTAGTAGGTCGTCTTG |
| mew_ex4 | CCAGCTTCTAGCATCCATGCCCTATAAGCGGCGAATTCAAGTCCGAAT |
| mew_ex5 | CCAGCTTCTAGCATCCATGCCCTATAAGCAAGATAGCTGTACTTGTCC |
| mew_ex6 | CCAGCTTCTAGCATCCATGCCCTATAAGAAAGAATCTGCCACCGGTCA |
| mew_ex7 | CCAGCTTCTAGCATCCATGCCCTATAAGCAGCCGCATATGACATGTGA |
| mew_ex8 | CCAGCTTCTAGCATCCATGCCCTATAAGTGTCGAAAATCACCACCTGT |
| mew_ex9 | CCAGCTTCTAGCATCCATGCCCTATAAGGGTATTGGATTATCGGTGGA |
| mew_ex10 | CCAGCTTCTAGCATCCATGCCCTATAAGGAACCGAATTGCTCACCATC |
| mew_ex11 | CCAGCTTCTAGCATCCATGCCCTATAAGTTGCCAGTTCATAGCCAAAA |
| mew_ex12 | CCAGCTTCTAGCATCCATGCCCTATAAGGTGATCTCCGTTTATATCGG |
| mew_ex13 | CCAGCTTCTAGCATCCATGCCCTATAAGGGAGCTGCCACTATTAAATC |
| mew_ex14 | CCAGCTTCTAGCATCCATGCCCTATAAGCCTTCCGTTTTTGTGAAGTA |
| mew_ex15 | CCAGCTTCTAGCATCCATGCCCTATAAGCTGGTACACATACACAGCTC |
| mew_ex16 | CCAGCTTCTAGCATCCATGCCCTATAAGTGGGCAACGTATCCTGGATA |
| mew_ex17 | CCAGCTTCTAGCATCCATGCCCTATAAGCGTCAACTTGAGTGTGTACT |
| mew_ex18 | CCAGCTTCTAGCATCCATGCCCTATAAGACCGAAACGACTCTCCAGAG |
| mew_ex19 | CCAGCTTCTAGCATCCATGCCCTATAAGGTTCAGATCTCCGATATTGG |
| mew_ex20 | CCAGCTTCTAGCATCCATGCCCTATAAGAGTAGATGTAGACTACGCCG |
| mew_ex21 | CCAGCTTCTAGCATCCATGCCCTATAAGAATTCAGGCCCTGACTCGAG |
| mew_ex22 | CCAGCTTCTAGCATCCATGCCCTATAAGTTGACCATTGGGTATGGTTC |
| mew_ex23 | CCAGCTTCTAGCATCCATGCCCTATAAGTAGAGATGCCAAAGGTACGA |
| mew_ex24 | CCAGCTTCTAGCATCCATGCCCTATAAGTATCATCCAAGTCCGTGTTG |
| mew_ex25 | CCAGCTTCTAGCATCCATGCCCTATAAGATTACCACATCCGGATAGGA |
| mew_ex26 | CCAGCTTCTAGCATCCATGCCCTATAAGTGCCGAGGAGTTGAATGCTC |
| mew_ex27 | CCAGCTTCTAGCATCCATGCCCTATAAGACACTCGTCTGAATGCTGAT |
| mew_ex28 | CCAGCTTCTAGCATCCATGCCCTATAAGCGTATTGGGATCCATGTTAT |
| mew_ex29 | CCAGCTTCTAGCATCCATGCCCTATAAGACATGTTAGATTACTGGCCG |
| mew_ex30 | CCAGCTTCTAGCATCCATGCCCTATAAGTGCAACATGCTCTGAACGTG |
| mew_ex31 | CCAGCTTCTAGCATCCATGCCCTATAAGCTTCTCATCGTACGGTTCAA |
| mew_ex32 | CCAGCTTCTAGCATCCATGCCCTATAAGCTATAAGCCAACCGGAGTTC |
| mew_ex33 | CCAGCTTCTAGCATCCATGCCCTATAAGTGATCGAAGGTTTCCGCTTC |
| mew_ex34 | CCAGCTTCTAGCATCCATGCCCTATAAGAGACGCGAGAGAATTTCTTT |
| mew_ex35 | CCAGCTTCTAGCATCCATGCCCTATAAGCTTATTTTCCCGATCGAAGA |
| mew_ex36 | CCAGCTTCTAGCATCCATGCCCTATAAGCACGCGACTGAGCACATTCG |
| mew_ex37 | CCAGCTTCTAGCATCCATGCCCTATAAGTTCCATTCGTATGCACTCTC |
| mew_ex38 | CCAGCTTCTAGCATCCATGCCCTATAAGGGGTATTGGCCTTGATGTAG |
| mew_ex39 | CCAGCTTCTAGCATCCATGCCCTATAAGAAACGCACGGGAGTCTGGAT |
| mew_ex40 | CCAGCTTCTAGCATCCATGCCCTATAAGCACCAGCGAGTACTTCAAGC |
| mew_ex41 | CCAGCTTCTAGCATCCATGCCCTATAAGAAAGCAGAATCCGCCAATGG |
| mew_ex42 | CCAGCTTCTAGCATCCATGCCCTATAAGATCCAATATCGGATTGAGGC |
| mew_ex43 | CCAGCTTCTAGCATCCATGCCCTATAAGTCGAAATCGACATGCGCCTG |
| mew_ex44 | CCAGCTTCTAGCATCCATGCCCTATAAGACAGTCTTTCTGGAAGGTGC |
| mew_ex45 | CCAGCTTCTAGCATCCATGCCCTATAAGTCTCACAGAGATCGTCATCG |
| mew_ex46 | CCAGCTTCTAGCATCCATGCCCTATAAGTCCACCCGGATAATCAGATT |
| mew_ex47 | CCAGCTTCTAGCATCCATGCCCTATAAGTGAGGACTCAGTGATGTTCG |
| mew_ex48 | CCAGCTTCTAGCATCCATGCCCTATAAGCAGTCCATAATCTTCGTGTT |
| sog_ex1 | CCAGCTTCTAGCATCCATGCCCTATAAGattgtaggtggtgtacatgg |
| sog_ex2 | CCAGCTTCTAGCATCCATGCCCTATAAGttgtggaacaggaaacgggc |
| sog_ex3 | CCAGCTTCTAGCATCCATGCCCTATAAGgcgatgaggtgtagaaggag |
| sog_ex4 | CCAGCTTCTAGCATCCATGCCCTATAAGataacacccgcatcatcaac |
| sog_ex5 | CCAGCTTCTAGCATCCATGCCCTATAAGtgatagacactgagagtgcc |
| sog_ex6 | CCAGCTTCTAGCATCCATGCCCTATAAGgcagaatgcgcttgtaatca |
| sog_ex7 | CCAGCTTCTAGCATCCATGCCCTATAAGaactgaacaactccgtctgc |
| sog_ex8 | CCAGCTTCTAGCATCCATGCCCTATAAGcattgaagaccagggtgaga |
| sog_ex9 | CCAGCTTCTAGCATCCATGCCCTATAAGgctcaattttcacactcagt |
| sog_ex10 | CCAGCTTCTAGCATCCATGCCCTATAAGcacacgtggaatctcatcga |
| sog_ex11 | CCAGCTTCTAGCATCCATGCCCTATAAGacgcgacatcagtcgaagat |
| sog_ex12 | CCAGCTTCTAGCATCCATGCCCTATAAGatccatcggtgttcaagtag |
| sog_ex13 | CCAGCTTCTAGCATCCATGCCCTATAAGcaaactgatgttgggcctat |
| sog_ex14 | CCAGCTTCTAGCATCCATGCCCTATAAGtggttgaagttgaagctcgg |
| sog_ex15 | CCAGCTTCTAGCATCCATGCCCTATAAGaacttctccacactaccaat |
| sog_ex16 | CCAGCTTCTAGCATCCATGCCCTATAAGattgcactcgttgtcaatgg |
| sog_ex17 | CCAGCTTCTAGCATCCATGCCCTATAAGcgttgaattcctcgagcaat |
| sog_ex18 | CCAGCTTCTAGCATCCATGCCCTATAAGaagaagccttccagatagga |
| sog_ex19 | CCAGCTTCTAGCATCCATGCCCTATAAGttggagtgcttggaatggac |
| sog_ex20 | CCAGCTTCTAGCATCCATGCCCTATAAGcgattcgttgtagaagcgtc |
| sog_ex21 | CCAGCTTCTAGCATCCATGCCCTATAAGcacatctgacaggaatcctg |
| sog_ex22 | CCAGCTTCTAGCATCCATGCCCTATAAGgaggattgcgctgaatagtc |
| sog_ex23 | CCAGCTTCTAGCATCCATGCCCTATAAGcttcttgtctggacgatagg |
| sog_ex24 | CCAGCTTCTAGCATCCATGCCCTATAAGtcttgtgtccattggaactg |
| gt_ex1 | CCTCCTAAGTTTCGAGCTGGACTCAGTGcgatgtcgaacgcaaactga |
| gt_ex2 | CCTCCTAAGTTTCGAGCTGGACTCAGTGatgttgctatatcttcacgc |
| gt_ex3 | CCTCCTAAGTTTCGAGCTGGACTCAGTGatagactggtgatctagctc |
| gt_ex4 | CCTCCTAAGTTTCGAGCTGGACTCAGTGtaatccaactgattgacgct |
| gt_ex5 | CCTCCTAAGTTTCGAGCTGGACTCAGTGtgttcggtgtatggtctctg |
| gt_ex6 | CCTCCTAAGTTTCGAGCTGGACTCAGTGcatgagtttctcgtgcatta |
| gt_ex7 | CCTCCTAAGTTTCGAGCTGGACTCAGTGcttgcgatcagtcttgagat |
| gt_ex8 | CCTCCTAAGTTTCGAGCTGGACTCAGTGtgaaccggcaattggctgtg |
| gt_ex9 | CCTCCTAAGTTTCGAGCTGGACTCAGTGcgtgtacagatccattttgg |
| gt_ex10 | CCTCCTAAGTTTCGAGCTGGACTCAGTGggaaagatccaggacctctg |
| gt_ex11 | CCTCCTAAGTTTCGAGCTGGACTCAGTGtgtctctacgctgtcacatc |
| gt_ex12 | CCTCCTAAGTTTCGAGCTGGACTCAGTGtgtttgatacggcgagggag |
| gt_ex13 | CCTCCTAAGTTTCGAGCTGGACTCAGTGaaccactgccgtagctatag |
| gt_ex14 | CCTCCTAAGTTTCGAGCTGGACTCAGTGggcatacagaagattgctgg |
| gt_ex15 | CCTCCTAAGTTTCGAGCTGGACTCAGTGaaagcgggatacagggaggc |
| gt_ex16 | CCTCCTAAGTTTCGAGCTGGACTCAGTGctgcttgatgttgctgtagt |
| gt_ex17 | CCTCCTAAGTTTCGAGCTGGACTCAGTGaaggttagcggttggtgtaa |
| gt_ex18 | CCTCCTAAGTTTCGAGCTGGACTCAGTGttctccagaacgtctatctg |
| gt_ex19 | CCTCCTAAGTTTCGAGCTGGACTCAGTGtgaagggacgggtgttcttt |
| gt_ex20 | CCTCCTAAGTTTCGAGCTGGACTCAGTGcagctatgaccaagggatcc |
| gt_ex21 | CCTCCTAAGTTTCGAGCTGGACTCAGTGacatcagtggctgcgaaatt |
| gt_ex22 | CCTCCTAAGTTTCGAGCTGGACTCAGTGcactcgcggattgtccaaaa |
| gt_ex23 | CCTCCTAAGTTTCGAGCTGGACTCAGTGcatttttgggttggttacgg |
| gt_ex24 | CCTCCTAAGTTTCGAGCTGGACTCAGTGtgacggatccacttcttgag |
| gt_ex25 | CCTCCTAAGTTTCGAGCTGGACTCAGTGtgttgttgtttgaggagctg |
| gt_ex26 | CCTCCTAAGTTTCGAGCTGGACTCAGTGgagtcacagtcgcttgactc |
| gt_ex27 | CCTCCTAAGTTTCGAGCTGGACTCAGTGcggacttgctctcgaagttg |
| gt_ex28 | CCTCCTAAGTTTCGAGCTGGACTCAGTGcagtagcgttagccaagttg |
| gt_ex29 | CCTCCTAAGTTTCGAGCTGGACTCAGTGtagtatgcggcatccttaac |
| gt_ex30 | CCTCCTAAGTTTCGAGCTGGACTCAGTGgcagcattgttcttgcgacg |
| gt_ex31 | CCTCCTAAGTTTCGAGCTGGACTCAGTGcgatctcatcctccttgatg |
| gt_ex32 | CCTCCTAAGTTTCGAGCTGGACTCAGTGtgttctggcgttccagatag |
| gt_ex33 | CCTCCTAAGTTTCGAGCTGGACTCAGTGtcgatctggcacaagagctc |
| gt_ex34 | CCTCCTAAGTTTCGAGCTGGACTCAGTGttactttggcggaggtgaag |
| gt_ex35 | CCTCCTAAGTTTCGAGCTGGACTCAGTGagagaggagtggacctttag |
| gt_ex36 | CCTCCTAAGTTTCGAGCTGGACTCAGTGattcggatcctcgcgttcaa |
| gt_ex37 | CCTCCTAAGTTTCGAGCTGGACTCAGTGgctgacccaaaaactggaca |
| gt_ex38 | CCTCCTAAGTTTCGAGCTGGACTCAGTGcccctaaactattcacagat |
| gt_ex39 | CCTCCTAAGTTTCGAGCTGGACTCAGTGtacagctaatacattttcca |
| gt_ex40 | CCTCCTAAGTTTCGAGCTGGACTCAGTGctggtgggattgcgagatgc |
| gt_ex41 | CCTCCTAAGTTTCGAGCTGGACTCAGTGagatgattccattgatagtt |
| gt_ex42 | CCTCCTAAGTTTCGAGCTGGACTCAGTGtagctgggacttctacaact |
| gt_ex43 | CCTCCTAAGTTTCGAGCTGGACTCAGTGtcattccttagggctatgaa |
